## Supplemental Figures and Tables for "Structural Basis for Sarbecovirus Rc-o319 Spike Adaptation to *Rhinolophus cornutus* Bat ACE2 and Constraints on Switching to Human ACE2"

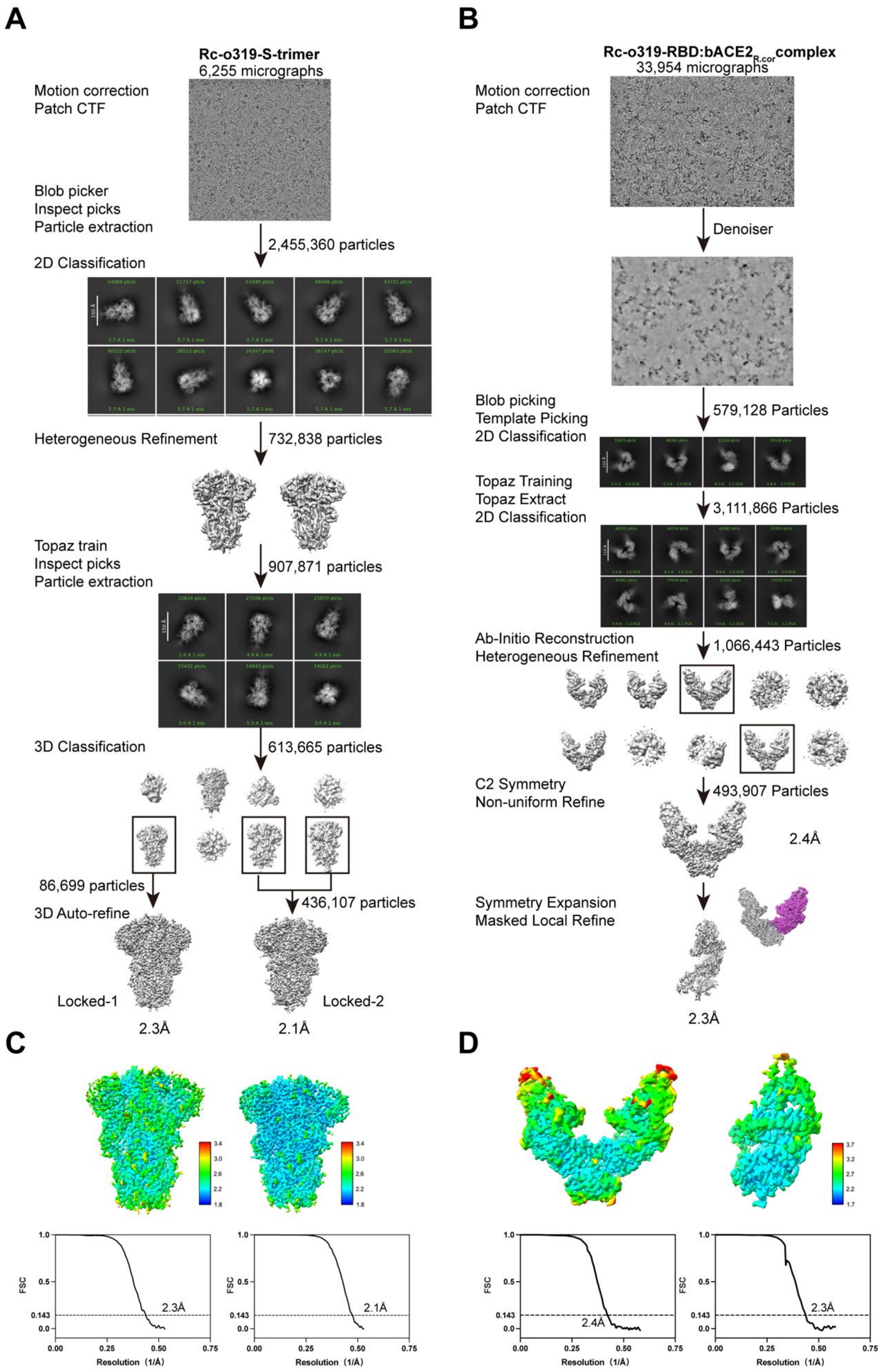
A
Rc-o319-S-trimer
6,255 micrographs
Motion correction
Patch CTF
Blob picker
Inspect picks
Particle extraction
2,455,360 particles
2D Classification
Heterogeneous Refinement
732,838 particles
Topaz train
Inspect picks
Particle extraction
907,871 particles
3D Classification
613,665 particles
86,699 particles
436,107 particles
3D Auto-refine
Locked-1
Locked-2
2.3Å
2.1Å
B
Rc-o319-RBD:bACE2R.cor complex
33,954 micrographs
Motion correction
Patch CTF
Denoiser
Blob picking
Template Picking
2D Classification
579,128 Particles
Topaz Training
Topaz Extract
2D Classification
3,111,866 Particles
Ab-Initio Reconstruction
Heterogeneous Refinement
1,066,443 Particles
C2 Symmetry
Non-uniform Refine
493,907 Particles
2.4Å
Symmetry Expansion
Masked Local Refine
2.3Å
C
FSC
Resolution (1/Å)
2.3Å
2.1Å
D
2.4Å
2.3Å

**Figure S1. Cryo-EM data processing and structure analysis for the Rc-o319 S-trimer and Rc-o319-RBD: bACE2$_{R.cor}$ complex datasets.**

**(A-B)** Data processing pipelines are shown for the Rc-o319 S-trimer locked-1 and locked-2 structures and Rc-o319-RBD:bACE2$_{R.cor}$ structures. 3D and 2D classification steps were used to remove contaminating particles. Two conformations were identified in the 3D classification for the Rc-o319 S-trimer.
**(C-D)** Local resolution maps for the Rc-o319 S-trimer and Rc-o319-RBD:bACE2$_{R.cor}$ complex structures (top panels) and global resolution assessments by Fourier shell correlation at the 0.143 criterion (bottom panels).

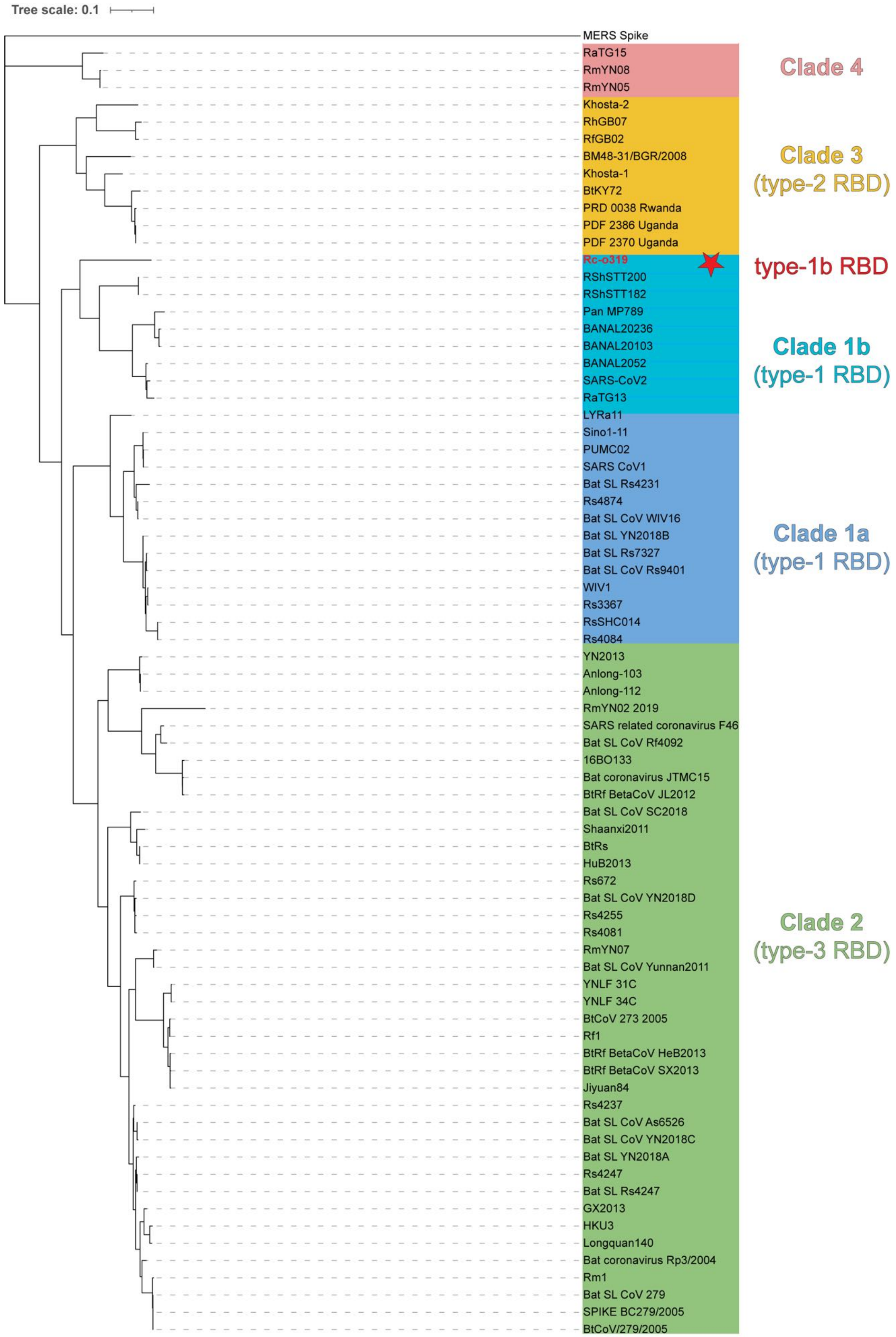

Tree scale: 0.1
MERS Spike
RaTG15
RmYN08
RmYN05
Khosta-2
RhGB07
RfGB02
BM48-31/BGR/2008
Khosta-1
BtKY72
PRD 0038 Rwanda
PDF 2386 Uganda
PDF 2370 Uganda
Rc-o319
RShSTT200
RShSTT182
Pan MP789
BANAL20236
BANAL20103
BANAL2052
SARS-CoV2
RaTG13
LYRa11
Sino1-11
PUMC02
SARS CoV1
Bat SL Rs4231
Rs4874
Bat SL CoV WIV16
Bat SL YN2018B
Bat SL Rs7327
Bat SL CoV Rs9401
WIV1
Rs3367
RsSHC014
Rs4084
YN2013
Anlong-103
Anlong-112
RmYN02 2019
SARS related coronavirus F46
Bat SL CoV Rf4092
16BO133
Bat coronavirus JTMC15
BtRf BetaCoV JL2012
Bat SL CoV SC2018
Shaanxi2011
BtRs
HuB2013
Rs672
Bat SL CoV YN2018D
Rs4255
Rs4081
RmYN07
Bat SL CoV Yunnan2011
YNLF 31C
YNLF 34C
BtCoV 273 2005
Rf1
BtRf BetaCoV HeB2013
BtRf BetaCoV SX2013
Jiyuan84
Rs4237
Bat SL CoV As6526
Bat SL CoV YN2018C
Bat SL YN2018A
Rs4247
Bat SL Rs4247
GX2013
HKU3
Longquan140
Bat coronavirus Rp3/2004
Rm1
Bat SL CoV 279
SPIKE BC279/2005
BtCoV/279/2005
Clade 4
Clade 3
(type-2 RBD)
type-1b RBD
Clade 1b
(type-1 RBD)
Clade 1a
(type-1 RBD)
Clade 2
(type-3 RBD)

**Figure S2. Phylogenetic tree constructed based on 72 sarbecovirus S-protein amino acid sequences, with MERS S-protein used as the outgroup. The tree shows the evolutionary relationships based on sequence similarities.**

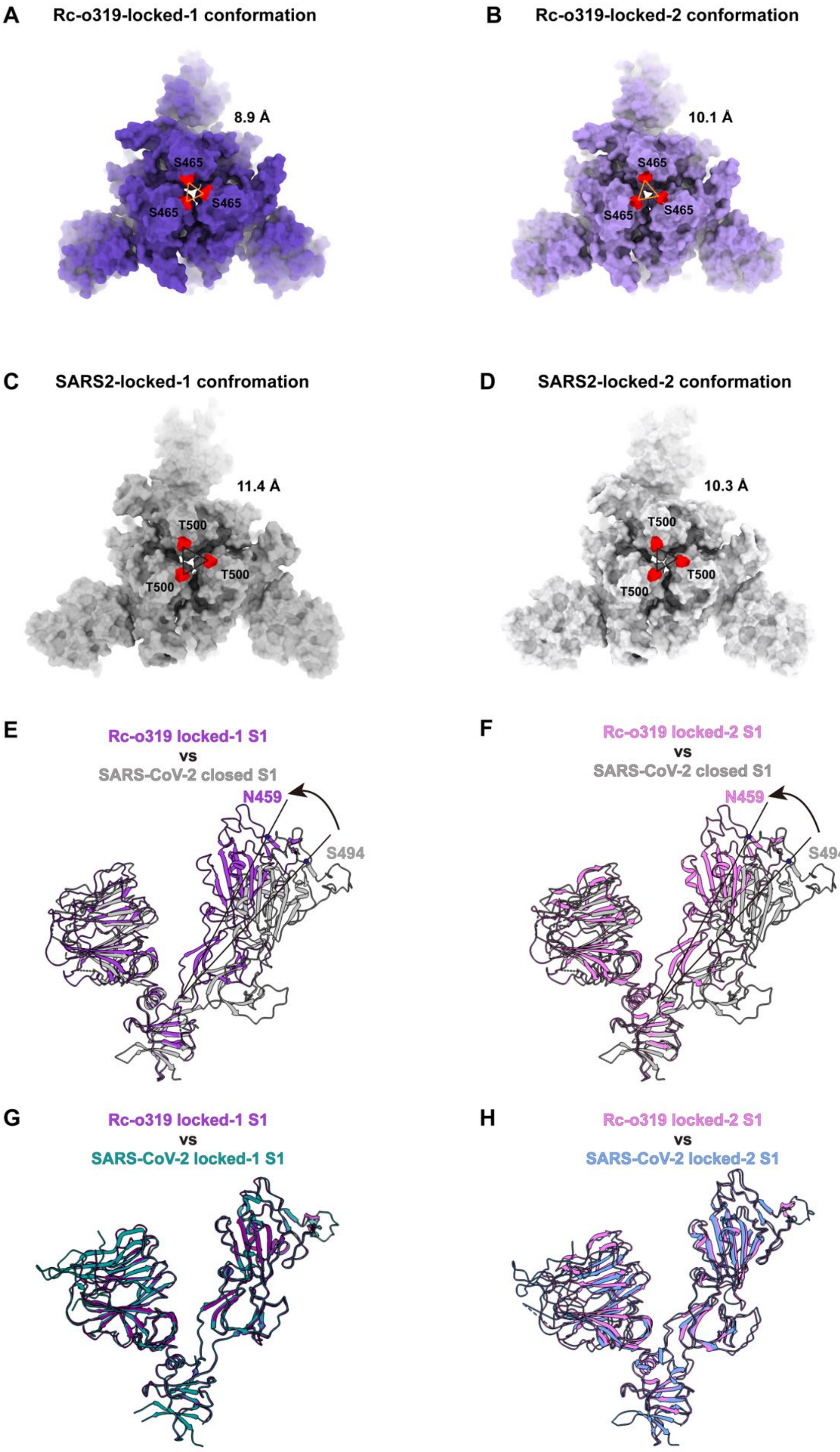
A
Rc-o319-locked-1 conformation
8.9 Å
S465
S465
S465
B
Rc-o319-locked-2 conformation
10.1 Å
S465
S465
S465
C
SARS2-locked-1 confromation
11.4 Å
T500
T500
T500
D
SARS2-locked-2 conformation
10.3 Å
T500
T500
T500
E
Rc-o319 locked-1 S1
vs
SARS-CoV-2 closed S1
N459
S494
F
Rc-o319 locked-2 S1
vs
SARS-CoV-2 closed S1
N459
S494
G
Rc-o319 locked-1 S1
vs
SARS-CoV-2 locked-1 S1
H
Rc-o319 locked-2 S1
vs
SARS-CoV-2 locked-2 S1

**Figure S3. Structural differences between Rc-o319 S-trimers in locked-1 and locked-2 conformations and comparison with SARS-CoV-2 S-trimer structures of different conformations.**

**(A-B)** Top-views of the two locked Rc-o319 S-trimer structures shown in molecular surface representation. Apex residues ($S465_{Rc\text{-}o319}$) in each S-trimer are colored red. In each S-trimer structure, the distance between apices is indicated by triangles, with the distances indicated. **(C-D)** Top-views of locked-1 (PDB: 7XTZ) and locked-2 (PDB: 7XU2) SARS-CoV-2 S-trimer structures, apex positions ($T500_{SARS2}$) are colored red, with the inter-apex distances indicated. **(E-F)** Comparison of the determined locked-1 and locked-2 Rc-o319 S1 structures (extracted from the Rc-o319 S-trimer structures) with the SARS-CoV-2 S1 structure in the closed conformation (gray, PDB: 7XU3). In the locked Rc-o319 S-trimers, the receptor-binding domains (RBDs) are positioned closer to the N-terminal domains (NTDs). (**G-H)** The S1 structures within the locked Rc-o319 S-trimers closely resemble those observed in the corresponding locked SARS-CoV-2 S-trimers (locked-1 PDB: 7XTZ, locked-2 PDB: 7XU2).

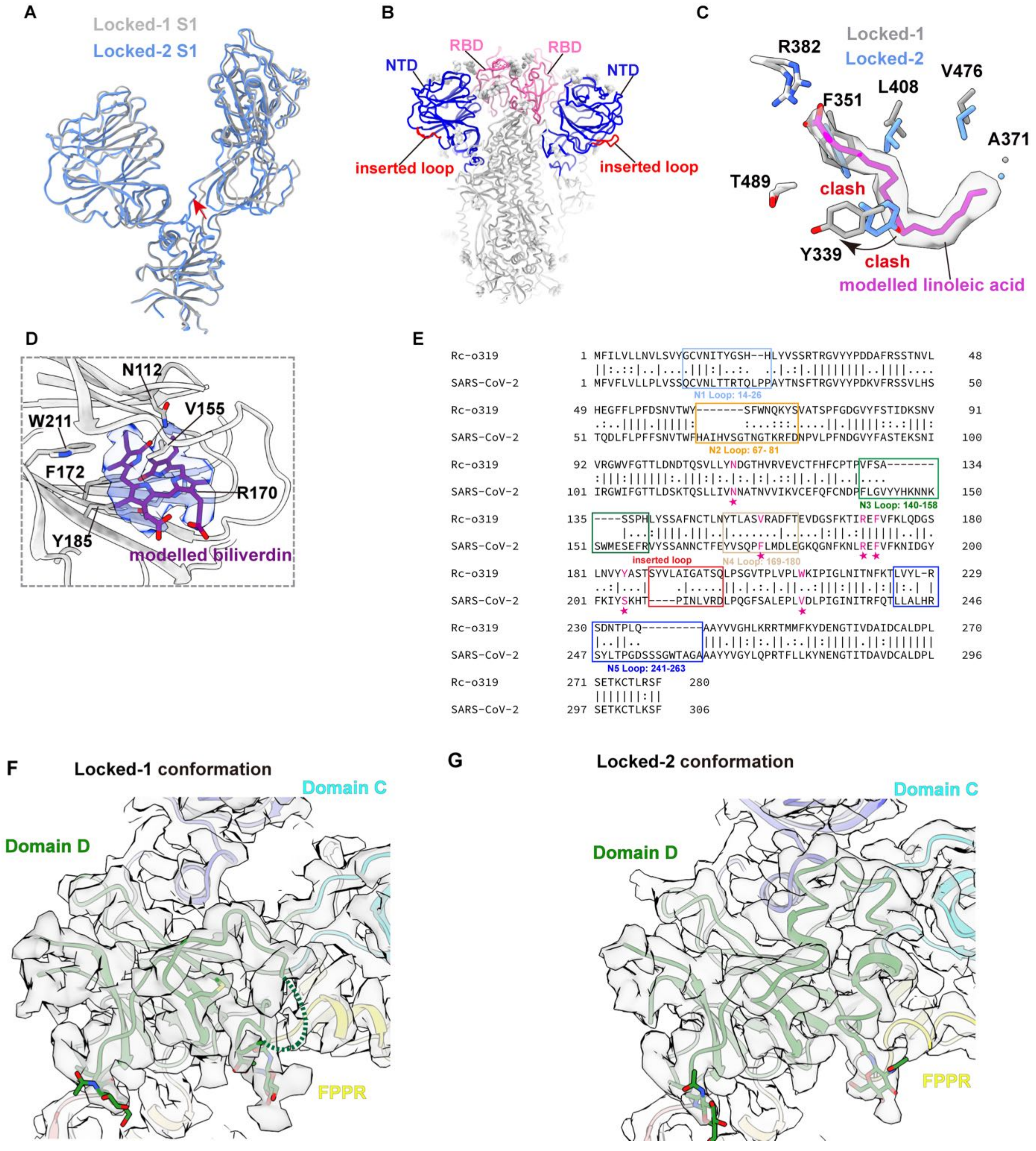


**Figure S4. Structural features of the Rc-o319 S-trimers in the two locked conformations.**

**(A)** Comparison of the locked-1 and locked-2 S1 structures of Rc-o319 S-trimer. **(B)** Rc-o319 locked-1 S-trimer structure showing the NTD and RBD. The "inserted loop" exposed to the exterior of the S-trimer is highlighted in red. **(C)** Comparison of the fatty-acid binding pockets of the fatty-acid-bound locked-1 and the fatty-acid-unbound locked-2 Rc-o319 S-trimer structures. A notable reorientation of the Y339$_{Rc\text{-}o319}$ side-chain (highlighted by red labels) is observed between the occupied and unoccupied fatty-acid binding pockets. Due to the contraction of the fatty-acid binding pocket, the side-chains of Y339$_{Rc\text{-}o319}$ and F351$_{Rc\text{-}o319}$ in the unoccupied fatty-acid pocket would clash with the modelled linoleic acid molecule, rendering the unoccupied pocket incompatible with lipid binding. **(D)** Modelled biliverdin is shown in purple stick representation with its density (blue). Residues involved in hydrophobic, hydrogen-bonding, and cation-π interactions with the bound biliverdin are shown as sticks. **(E)** An alignment of Rc-o319 and SARS-CoV-2 N-terminal domain (NTD) amino acid sequences. Modelled biliverdin interacting residues are marked by purple stars in the alignment. N1-N5 loops are framed by boxes colored in light blue, orange, green, brown and blue, respectively. Additionally, a four-amino-acid insertion, forming the "inserted loop" in the Rc-o319 NTD by comparison with SARS-CoV-2 NTD, is framed by a red box. **(F-G)** Cryo-EM densities of Domain D and the surrounding regions in locked-1 and locked-2 conformations. In the locked-1 conformation, Domain D contains a large disordered Domain D-loop; In locked-2 conformation, Domain D is fully ordered, with the disordered Domain D-loop in locked-1 refolded into two short α-helices.

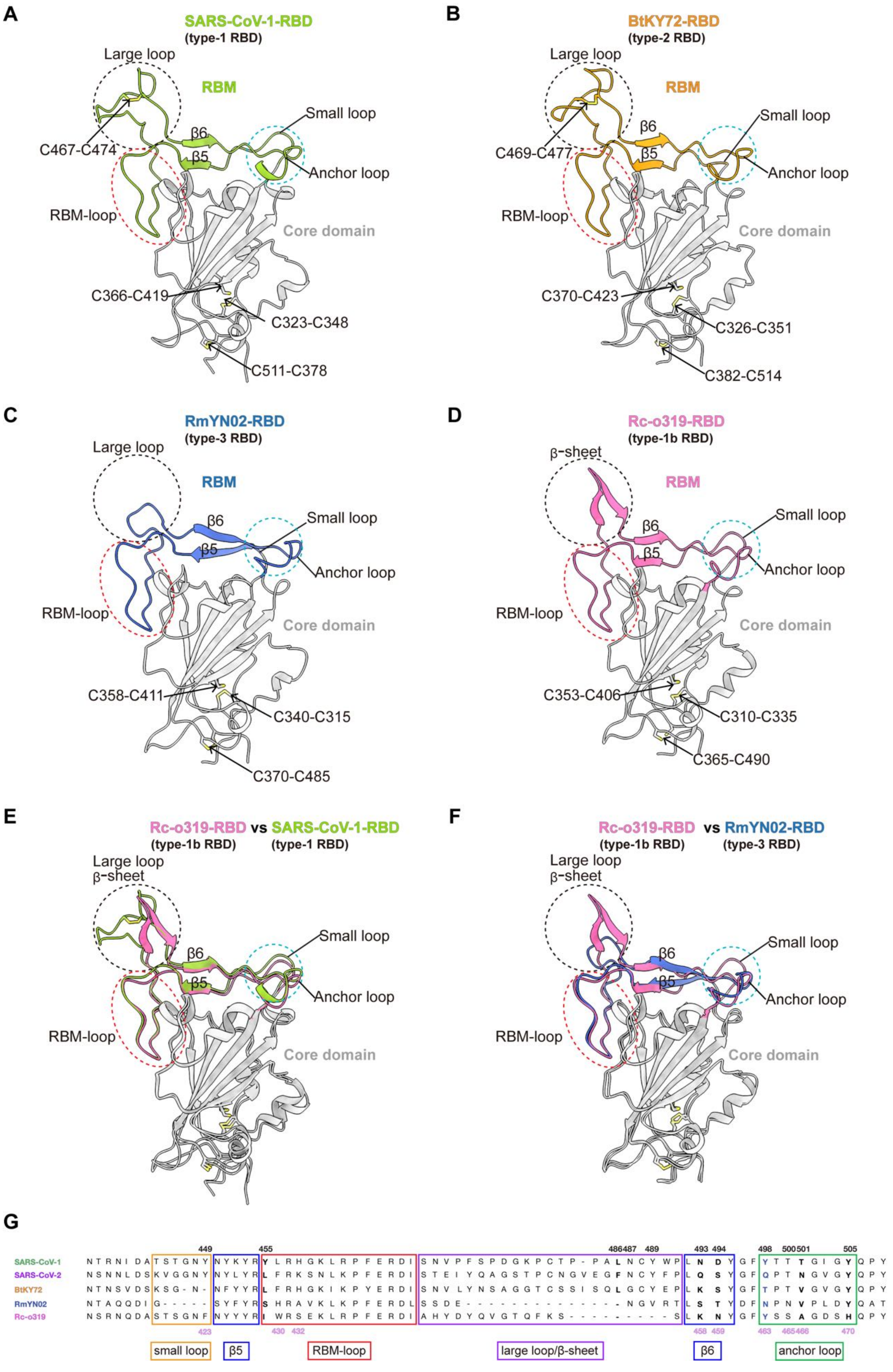
A
SARS-CoV-1-RBD
(type-1 RBD)
Large loop
RBM
C467-C474
β6
β5
Small loop
Anchor loop
RBM-loop
Core domain
C366-C419
C323-C348
C511-C378
B
BtKY72-RBD
(type-2 RBD)
Large loop
RBM
C469-C477
β6
β5
Small loop
Anchor loop
RBM-loop
Core domain
C370-C423
C326-C351
C382-C514
C
RmYN02-RBD
(type-3 RBD)
Large loop
RBM
β6
β5
Small loop
Anchor loop
RBM-loop
Core domain
C358-C411
C340-C315
C370-C485
D
Rc-o319-RBD
(type-1b RBD)
β-sheet
RBM
β6
β5
Small loop
Anchor loop
RBM-loop
Core domain
C353-C406
C310-C335
C365-C490
E
Rc-o319-RBD vs SARS-CoV-1-RBD
(type-1b RBD) (type-1 RBD)
Large loop
β-sheet
β6
β5
Small loop
Anchor loop
RBM-loop
Core domain
F
Rc-o319-RBD vs RmYN02-RBD
(type-1b RBD) (type-3 RBD)
Large loop
β-sheet
β6
β5
Small loop
Anchor loop
RBM-loop
Core domain
G
449 455 486 487 489 493 494 498 500 501 505
SARS-CoV-1 N T R N I D A T S T G N Y N Y K Y R Y L R H G K L R P F E R D I S N V P F S P D G K P C T P - P A L N C Y W P L N D Y G F Y T T T G I G Y Q P Y
SARS-CoV-2 N S N N L D S K V G G N Y N Y L Y R L F R K S N L K P F E R D I S T E I Y Q A G S T P C N G V E G F N C Y F P L Q S Y G F Q P T N G V G Y Q P Y
BtKY72 N T N S V D S K S G - N - N F Y Y R L F R H G K I K P Y E R D I S N V L Y N S A G G T C S S I S Q L G C Y E P L K S Y G F T P T V G V G Y Q P Y
RmYN02 N T A Q Q D I G - - - - - S Y F Y R S H R A V K L K P F E R D L S S D E - - - - - - - - - - - - - - N G V R T L S T Y D F N P N V P L D Y Q A T
Rc-o319 N S R N Q D A S T S G N F N Y Y Y R I W R S E K L R P F E R D I A H Y D Y Q V G T Q F K S - - - - - - - - - - S L K N Y G F Y S S A G D S H Q P Y
423 430 432 458 459 463 465 466 470
small loop
β5
RBM-loop
large loop/β-sheet
β6
anchor loop

**Figure S5. Representative type-1, type-2, type-3, and type-1b sarbecovirus RBD structures with key features highlighted.**

**(A-D)** Structures of representative type-1, type-2, type-3, and type-1b RBDs are shown in the same orientation. The RBM (receptor binding motif) regions are highlighted in colors, while the conserved RBD core domains are colored in gray. Black dashed circles are shown to mark the locations of the SL (Small-loop), AL (anchor-loop), and LL (large-loop) or beta-loop to highlight structural differences in the RBMs of different types of RBDs. Red dashed ovals highlight the RBM-loop structures. **(E)** Superimposition of the type-1b Rc-o319-RBD and the type-1 SARS-CoV-1-RBD structures. **(F)** Superimposition of the type-1b Rc-o319-RBD and the type-3 RmYN02-RBD structures. **(G)** An alignment of representative types 1-3 and type-1b RBM amino acid sequences. Amino acid residue numbers are shown according to SARS-CoV-2 (black, top) and Rc-o319 (pink, bottom) sequences.

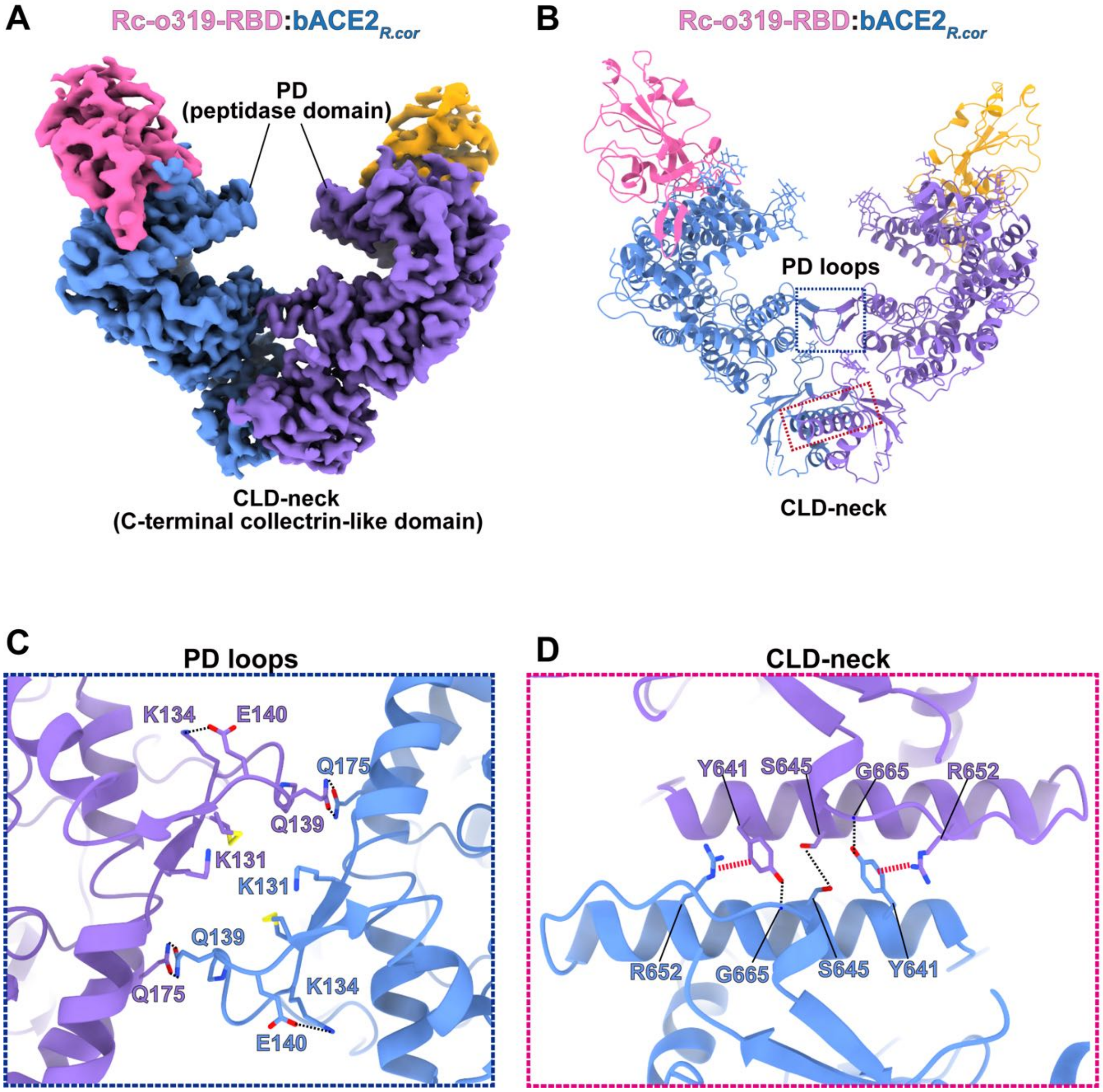


**Figure S6. Cryo-EM density and the associated molecular model of the dimeric Rc-o319-RBD:bACE2$_{R.cor}$ complex.**

**(A-B)** Cryo-EM density (A) and the associated molecular model (B) of the Rc-o319-RBD:bACE2$_{R.cor}$ dimer. **(C)** Detailed dimer interface interactions in the peptidase domain (PD) loop region. (**D)** Detailed dimer interface interactions in the C-terminal collectrin-like domain (CLD) neck region. Hydrogen bonds are shown as black dashed lines, salt bridges are shown as blue dashed lines, and cation-π interactions are shown as red dashed lines. Dashed boxes in (B) indicate the locations of the interfaces shown in panels C and D.

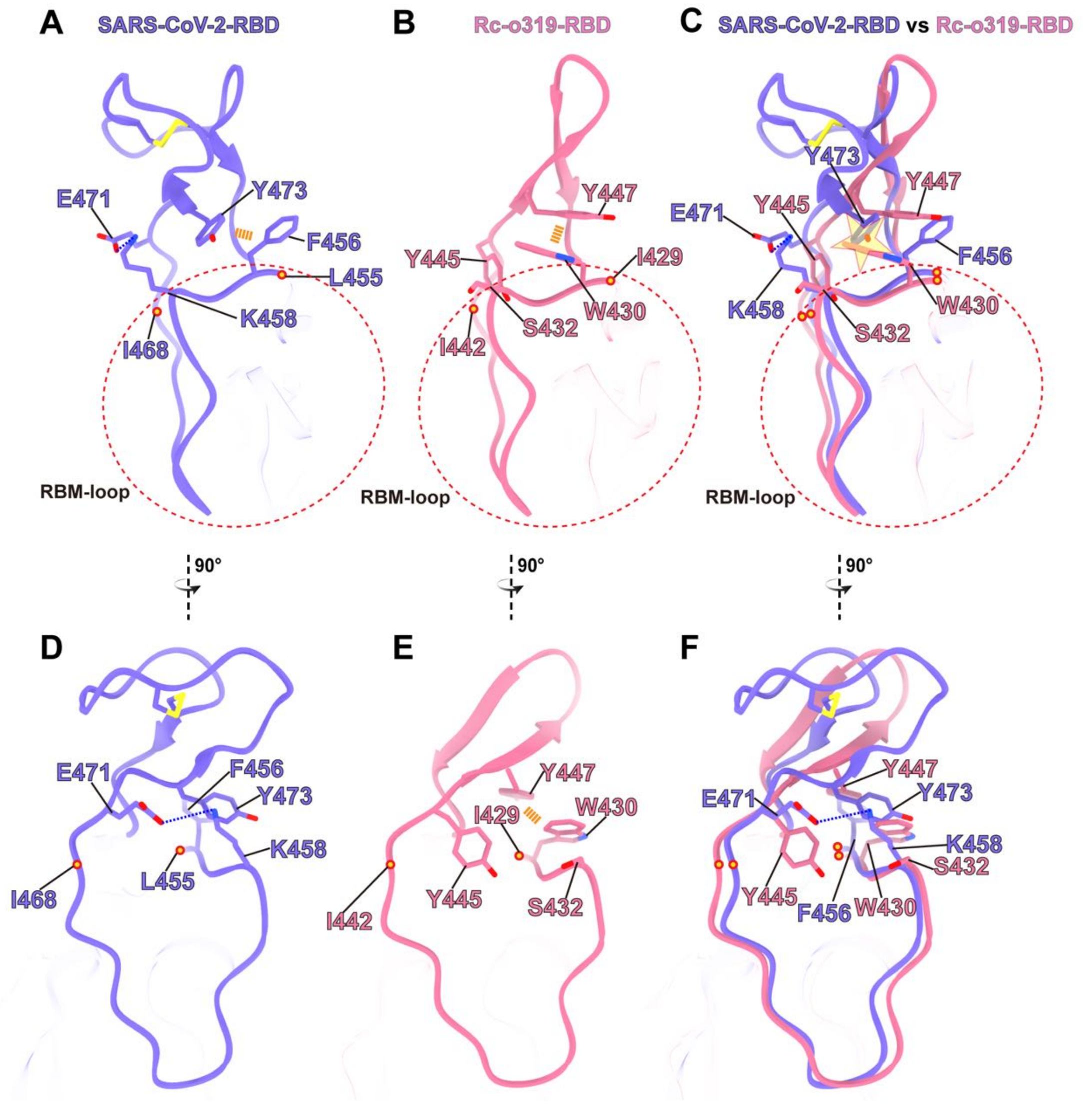


**Figure S7. Interactions between the RBM-loop and the beta-loop or the large-loop of the Rc-o319 and SARS-CoV-2 RBDs.**

**(A-B** and **D-E)** The beta-loop-RBM-loop region of Rc-o319-RBD and the large-loop-RBM-loop region of SARS-CoV-2-RBD are shown in two different viewing angles. **(A)** $Y473_{SARS2}$ of the SARS-CoV-2 large-loop and $F456_{SARS2}$ of the RBM-loop form a pi-pi interaction. **(B)** The corresponding residues $Y447_{Rc\text{-}o319}$ of the Rc-o319 beta-loop and $W430_{Rc\text{-}o319}$ of the Rc-o319 RBM-loop also form a pi-pi interaction in a substantially different conformation. **(C)** A superposition of the Rc-o319 beta-loop-RBM-loop and the SARS-CoV-2 large-loop-RBM-loop structures. The superposition reveals a clash (highlighted by a yellow star) between the side-chains of $Y473_{SARS2}$ and $W430_{Rc\text{-}o319}$. **(D)** In the rotated view, $E471_{SARS2}$ forms a salt-bridge with the RBM-loop residue $K458_{SARS2}$, likely stabilizing the SARS-

CoV-2 large-loop. **(E)** The corresponding residues in Rc-o319, $Y445_{Rc\text{-}o319}$ of beta-loop and $S432_{Rc\text{-}o319}$ of RBM-loop are not interacting. **(F)** The superposition of the Rc-o319 beta-loop-RBM-loop and the SARS-CoV-2 large-loop-RBM-loop structures in a rotated view.

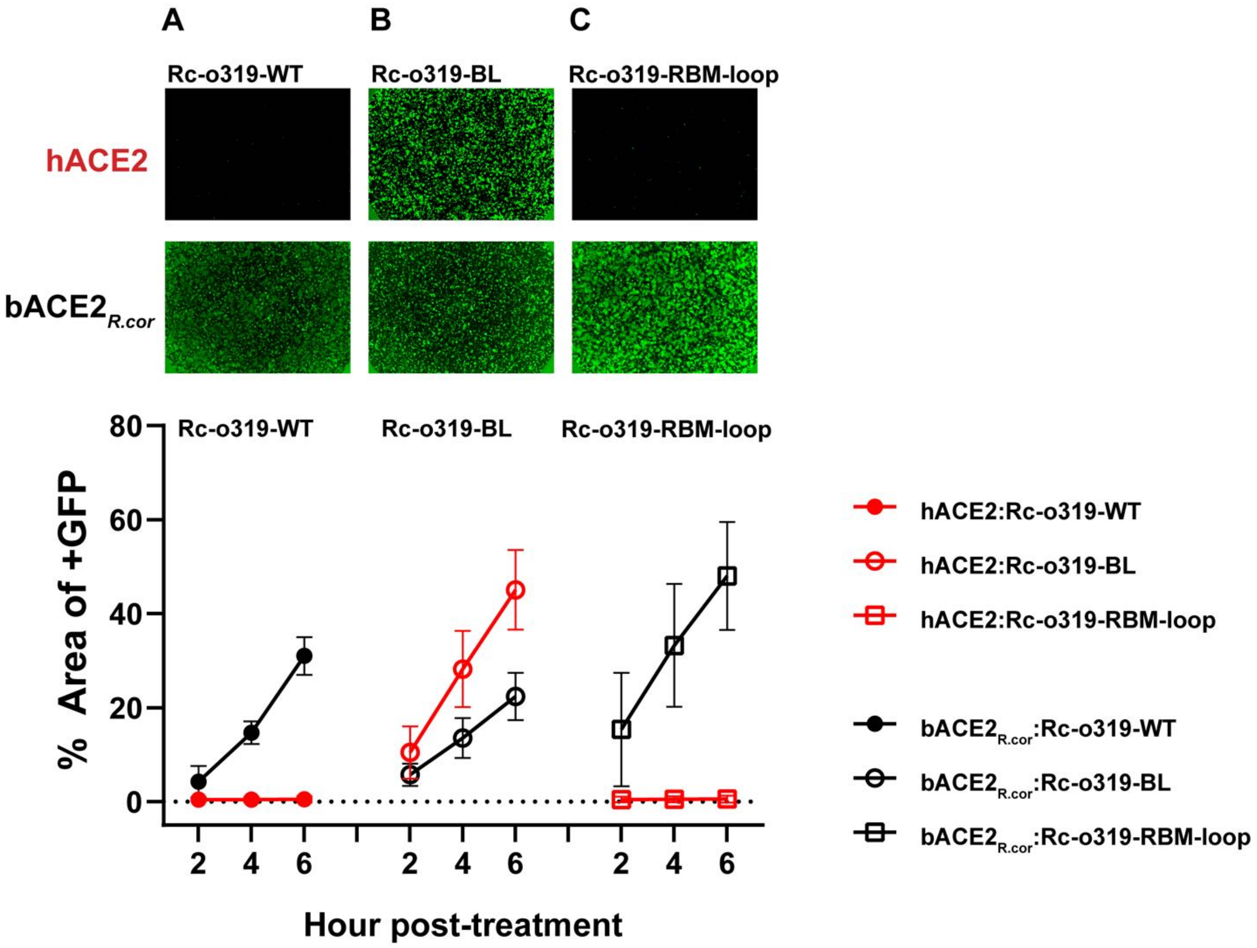


**Figure S8. ACE2 utilization by wild-type (WT) and variant Rc-o319 S proteins in cell-cell fusion assays.**

**(A-C)** Representative cell-cell fusion images captured at 6 hours post-transfection are shown. Effector cells expressing WT and variant Rc-o319 S-proteins were tested against receptor cells expressing either bACE2*R.cor* or hACE2. The Bottom panel: Cell-cell fusion was quantified by assessing GFP+ areas at 2, 4, and 6 hours post-transfection.

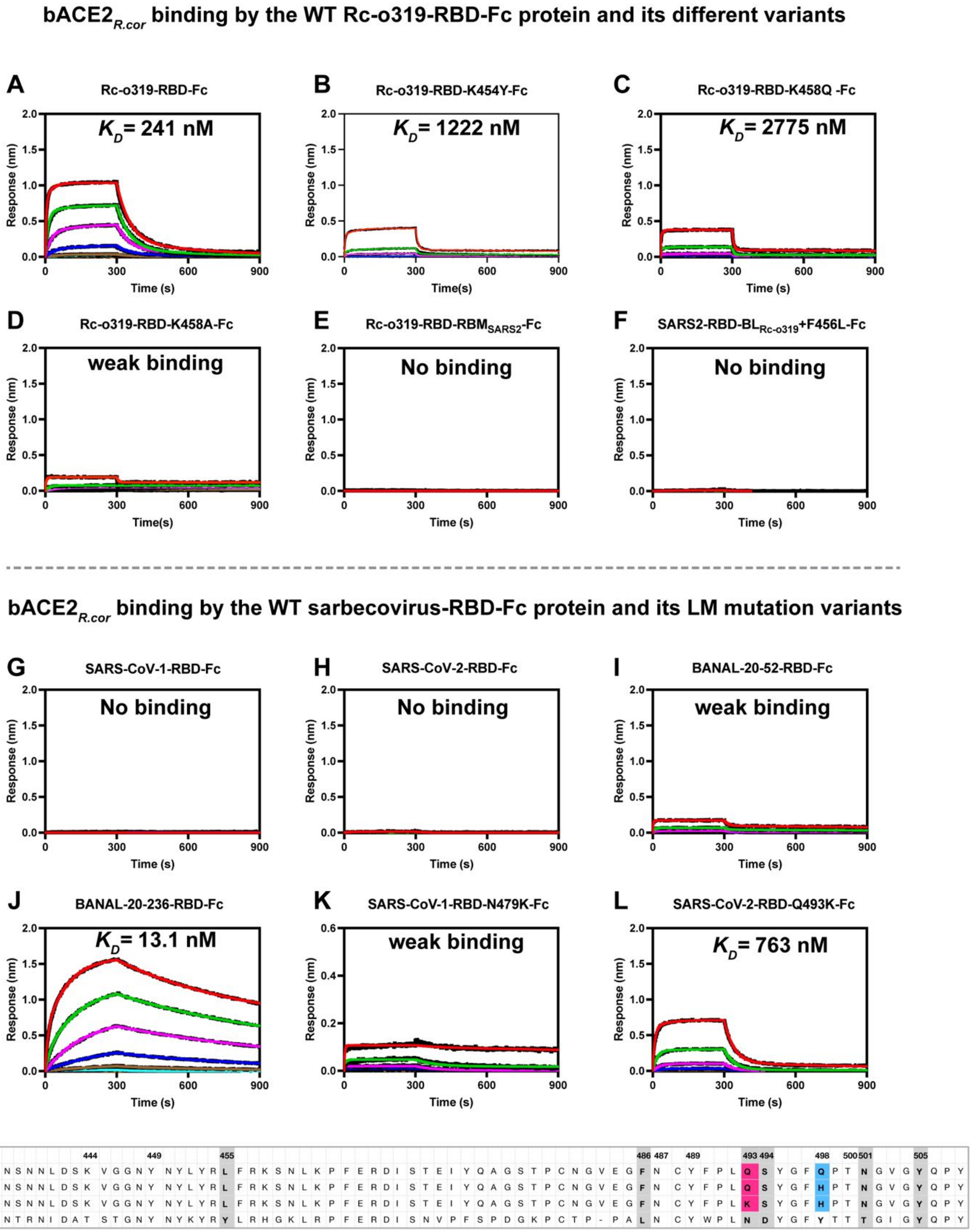


**Figure S9. Binding of bACE2*$_{R.cor}$* by wild-type (WT) and variant Rc-o319-RBD-Fc proteins, compared with RBD-Fc proteins from SARS-CoV-1, SARS-CoV-2, BANAL-20-52 and BANAL-20-236.**

**(A-E)** bACE2$_{R.cor}$ binding by the WT Rc-o319-RBD-Fc protein and its different variants, including K454Y$_{Rc\text{-}o319}$ in BL region (B) K458Q/A$_{Rc\text{-}o319}$ in LM (C and D), Rc-o319 RBD RBM exchanged for SARS-CoV-2 RBM (RBM$_{SARS2}$) (E) and **(F)** SARS-CoV-2-RBD-Fc with LL exchanged for BL of Rc-o319 (BL$_{Rc\text{-}o319}$) with an extra F456L$_{Rc\text{-}o319}$ mutation in the RBM-loop. **(G-J)** bACE2$_{R.cor}$ binding by wild-type RBD-Fc proteins of SARS-CoV-1, SARS-CoV-2, BANAL-20-52, and BANAL-20-236. (**K-L)** bACE2$_{R.cor}$ binding by LM variants of SARS-CoV-1-RBD-Fc (N479K$_{SARS1}$) and SARS-CoV-2-RBD-Fc (Q493K$_{SARS2}$) proteins. **(M)** An alignment of SARS-CoV-2, BANAL-20-236, BANAL-20-52, and SARS-CoV-1 RBM amino acid sequences. Compared to the SARS-CoV-2 RBM, there are two amino-acid changes, Q493K$_{SARS2}$ and Q498H$_{SARS2}$, in BANAL-20-236 and one amino-acid change, Q498H$_{SARS2}$, in BANAL-20-52. These changes likely favor bACE2$_{R.cor}$ binding by comparison with the SARS-CoV-2 Q493$_{SARS2}$ and Q498$_{SARS2}$ residues (H-J).

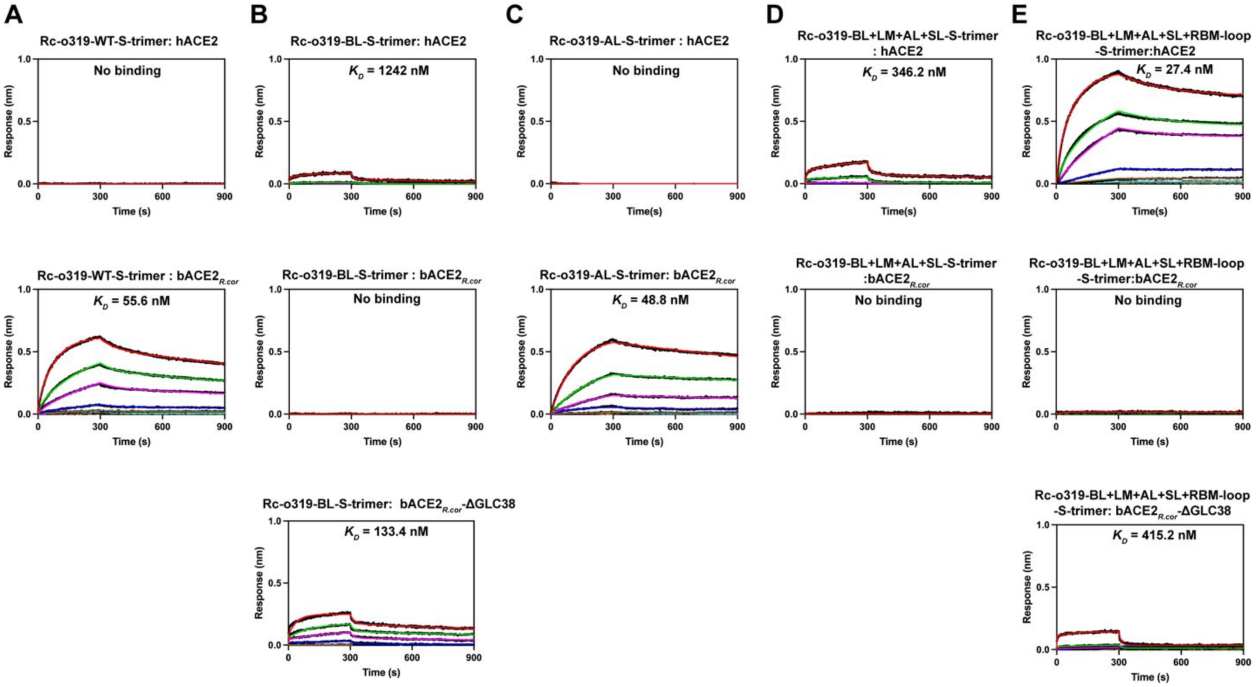


**Figure S10. Binding of wild-type (WT) bACE2*R.cor*, its Thr40Ala (removal of Asn38-glycan) variant (bACE2*R.cor*-ΔGLC38) and hACE2 by S-trimers of different Rc-o319 variants in BLI assays.**

**(A-E)** Binding sensorgrams were recorded by immersing biosensors immobilized with hACE2 (first panels), WT bACE2*R.cor* (second panels), or bACE2*R.cor*-ΔGLC38 (third panels) into three-fold serial dilutions of S-trimer solutions, with concentrations ranging from 3000 to 4.09 nM. For WT (A) and the Rc-o319 AL S protein (C), concentrations ranged from 1500 to 2.04 nM.

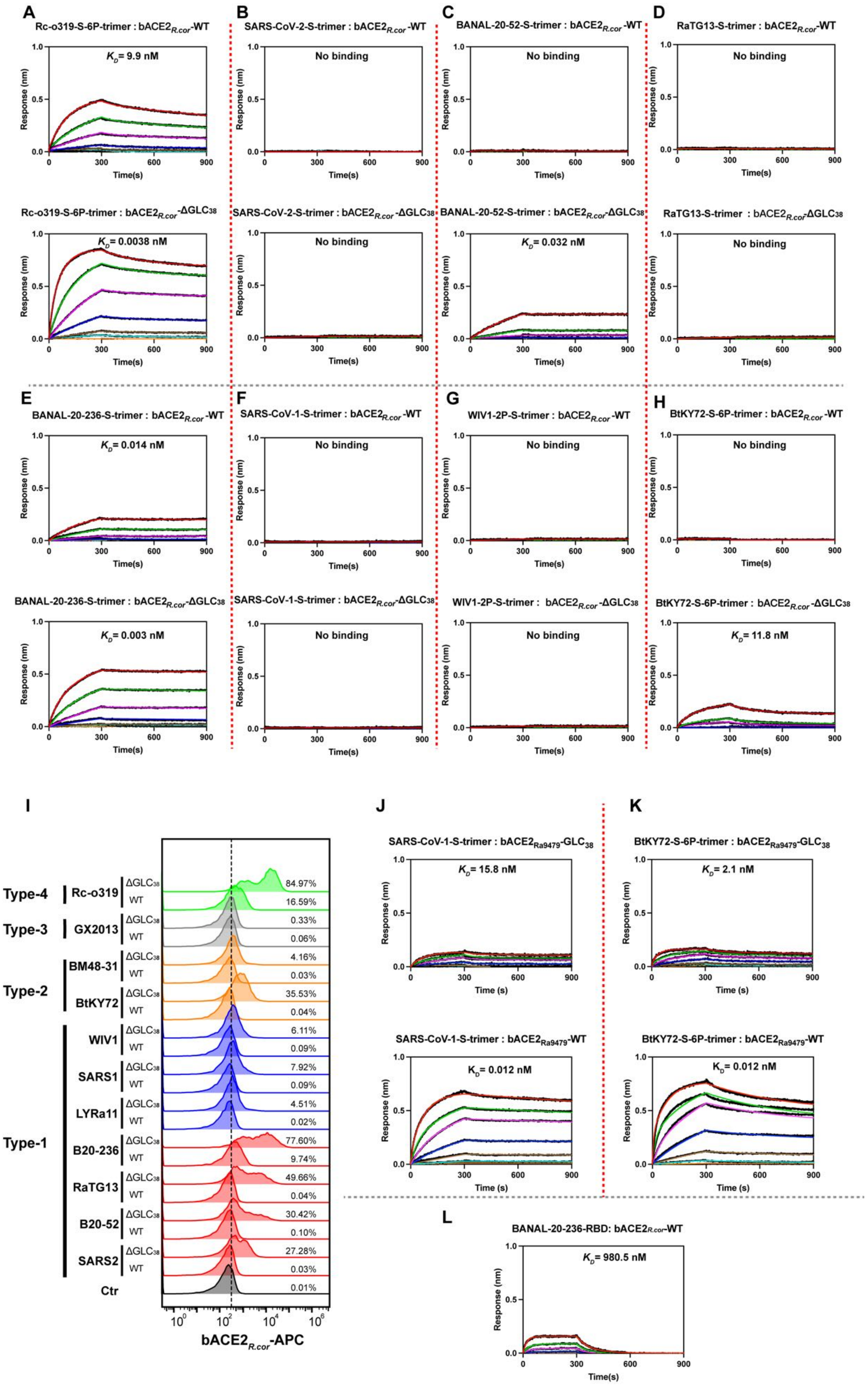
A
Rc-o319-S-6P-trimer : bACE2R.cor-WT
KD= 9.9 nM
Rc-o319-S-6P-trimer : bACE2R.cor-ΔGLC38
KD= 0.0038 nM
B
SARS-CoV-2-S-trimer : bACE2R.cor-WT
No binding
SARS-CoV-2-S-trimer : bACE2R.cor-ΔGLC38
No binding
C
BANAL-20-52-S-trimer : bACE2R.cor-WT
No binding
BANAL-20-52-S-trimer : bACE2R.cor-ΔGLC38
KD= 0.032 nM
D
RaTG13-S-trimer : bACE2R.cor-WT
No binding
RaTG13-S-trimer : bACE2R.cor-ΔGLC38
No binding
E
BANAL-20-236-S-trimer : bACE2R.cor-WT
KD= 0.014 nM
BANAL-20-236-S-trimer : bACE2R.cor-ΔGLC38
KD= 0.003 nM
F
SARS-CoV-1-S-trimer : bACE2R.cor-WT
No binding
SARS-CoV-1-S-trimer : bACE2R.cor-ΔGLC38
No binding
G
WIV1-2P-S-trimer : bACE2R.cor-WT
No binding
WIV1-2P-S-trimer : bACE2R.cor-ΔGLC38
No binding
H
BtKY72-S-6P-trimer : bACE2R.cor-WT
No binding
BtKY72-S-6P-trimer : bACE2R.cor-ΔGLC38
KD= 11.8 nM
Response (nm)
Time(s)
I
Type-4
Rc-o319
ΔGLC38 84.97%
WT 16.59%
Type-3
GX2013
ΔGLC38 0.33%
WT 0.06%
Type-2
BM48-31
ΔGLC38 4.16%
WT 0.03%
BtKY72
ΔGLC38 35.53%
WT 0.04%
Type-1
WIV1
ΔGLC38 6.11%
WT 0.09%
SARS1
ΔGLC38 7.92%
WT 0.09%
LYRa11
ΔGLC38 4.51%
WT 0.02%
B20-236
ΔGLC38 77.60%
WT 9.74%
RaTG13
ΔGLC38 49.66%
WT 0.04%
B20-52
ΔGLC38 30.42%
WT 0.10%
SARS2
ΔGLC38 27.28%
WT 0.03%
Ctr 0.01%
bACE2R.cor-APC
J
SARS-CoV-1-S-trimer : bACE2Ra9479-GLC38
KD= 15.8 nM
SARS-CoV-1-S-trimer : bACE2Ra9479-WT
KD= 0.012 nM
K
BtKY72-S-6P-trimer : bACE2Ra9479-GLC38
KD= 2.1 nM
BtKY72-S-6P-trimer : bACE2Ra9479-WT
KD= 0.012 nM
Time (s)
L
BANAL-20-236-RBD: bACE2R.cor-WT
KD= 980.5 nM

**Figure S11. Binding of wild-type (WT) bACE2$_{R.cor}$ and its Thr40Ala (removal of Asn38-glycan) variant (bACE2$_{R.cor}$-ΔGLC$_{38}$), by S-trimers of different sarbecoviruses in BLI and FACS assays.**

(**A-H**) Binding sensorgrams were recorded by submerging WT bACE2$_{R.cor}$ (top panels) or bACE2$_{R.cor}$-ΔGLC$_{38}$ (removing the glycan of Asn38 by Thr40Ala mutation, bottom panels) immobilized biosensors into 3-fold serially diluted S-trimer solutions, with concentrations ranging from 800 to 1.09 nM. **(I)** Binding of bACE2$_{R.cor}$-WT or bACE2$_{R.cor}$-ΔGLC$_{38}$ by cell-surface expressed S-proteins as assessed by flow cytometry. bACE2$_{R.cor}$-WT-Fc or bACE2$_{R.cor}$-ΔGLC$_{38}$-Fc protein was incubated with cells expressing S-proteins before ACE2 binding was quantified using a goat anti-human IgG-APC as the probe. (**J-K)** Binding of bACE2$_{Ra9479}$-WT and bACE2$_{Ra9479}$-GLC$_{38}$ (introducing the Asn38-glycan by the Thr40Ala mutation) by S-trimers of SARS-CoV-1 and BtKY72 as assessed by BLI assays. (**L**) Binding of bACE2$_{R.cor}$-WT by monomeric BANAL-20-236 RBD in BLI assays. Monomeric BANAL-20-236 RBD was threefold serially diluted from 3000 to 4.09 nM. Estimated $K_D$ values are shown next to their corresponding binding curves.

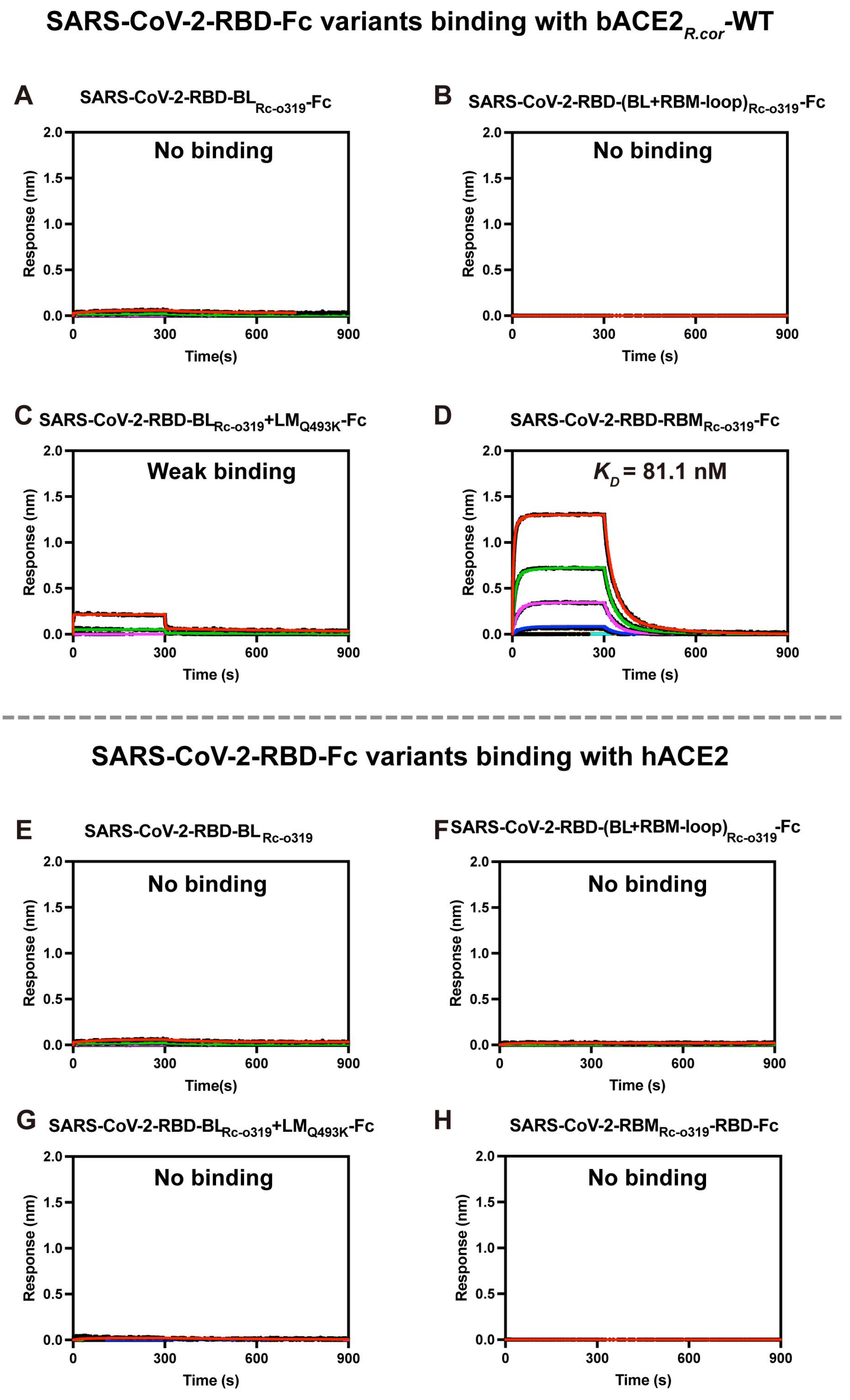

SARS-CoV-2-RBD-Fc variants binding with bACE2R.cor-WT
A
SARS-CoV-2-RBD-BLRc-o319-Fc
No binding
B
SARS-CoV-2-RBD-(BL+RBM-loop)Rc-o319-Fc
No binding
C
SARS-CoV-2-RBD-BLRc-o319+LMQ493K-Fc
Weak binding
D
SARS-CoV-2-RBD-RBMRc-o319-Fc
KD = 81.1 nM
Response (nm)
Time(s)
Time (s)
0
300
600
900
0.0
0.5
1.0
1.5
2.0
SARS-CoV-2-RBD-Fc variants binding with hACE2
E
SARS-CoV-2-RBD-BLRc-o319
No binding
F
SARS-CoV-2-RBD-(BL+RBM-loop)Rc-o319-Fc
No binding
G
SARS-CoV-2-RBD-BLRc-o319+LMQ493K-Fc
No binding
H
SARS-CoV-2-RBMRc-o319-RBD-Fc
No binding

**Figure S12. Binding of bACE2$_{R.cor}$ and hACE2 by SARS-CoV-2-RBD-Fc variant proteins incorporating Rc-o319 beta-loop and RBM-loop.**

**(A-D)** bACE2$_{R.cor}$ binding by different variants of SARS-CoV-2-RBD-Fc, including LL swapped for the Rc-o319 BL region (SARS-CoV-2-RBD-BL$_{Rc\text{-}o319}$-Fc) (A), swapped for the Rc-o319 BL region and RBM-loop (B), LL swapped for the Rc-o319 BL region plus lamella Q493$_{SARS2}$K mutation (SARS-CoV-2-RBD-BL$_{Rc\text{-}o319}$+LM$_{Q493K}$-Fc ) (C), and the whole SARS-CoV-2 RBM swapped for the Rc-o319 RBM (SARS-CoV-2-RBD-RBM$_{Rc\text{-}o319}$-Fc) (D). **(E-H)** hACE2 binding by SARS-CoV-2-RBD-Fc variants, corresponding to those in A-D. Dimeric SARS-CoV-2-RBD-Fc proteins were immobilized on the BLI sensors and tested binding against dimeric *Rhinolophus cornutus* bat ACE2 (bACE2$_{R.cor}$) or hACE2 protein as the analyte in solution. bACE2$_{R.cor}$ and hACE2 were threefold serially diluted from 3000 to 4.09 nM. Estimated $K_D$ values are shown next to their corresponding binding curves.

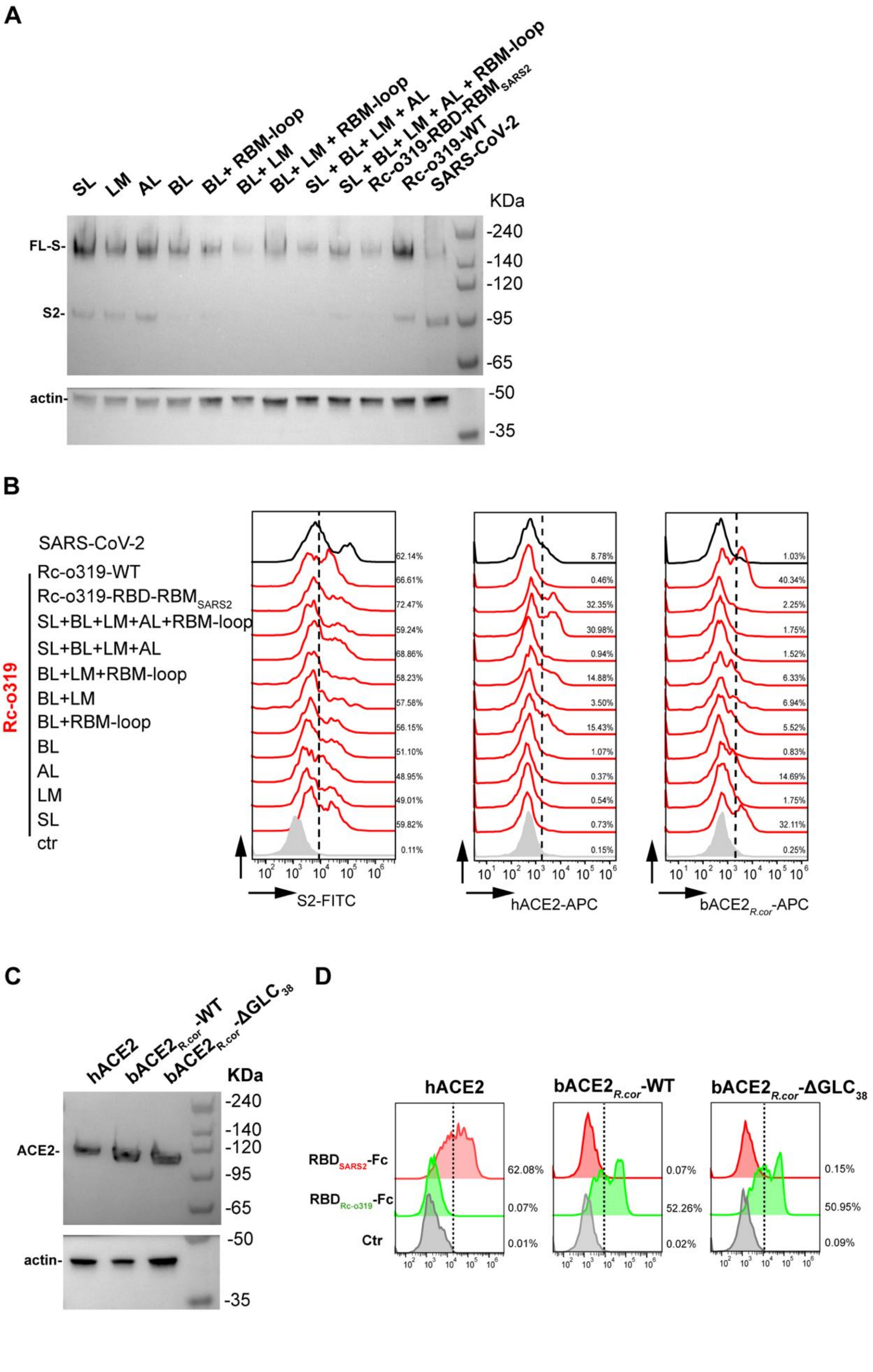

A
SL
LM
AL
BL
BL+ RBM-loop
BL+ LM
BL+ LM + RBM-loop
SL + BL+ LM + AL
SL + BL+ LM + AL + RBM-loop
Rc-o319-RBD-RBM$_{SARS2}$
Rc-o319-WT
SARS-CoV-2
KDa
FL-S-
S2-
actin-
-240
-140
-120
-95
-65
-50
-35
B
SARS-CoV-2
Rc-o319-WT
Rc-o319-RBD-RBM$_{SARS2}$
SL+BL+LM+AL+RBM-loop
SL+BL+LM+AL
BL+LM+RBM-loop
BL+LM
BL+RBM-loop
BL
AL
LM
SL
ctr
Rc-o319
62.14%
66.61%
72.47%
59.24%
68.86%
58.23%
57.58%
56.15%
51.10%
48.95%
49.01%
59.82%
0.11%
8.78%
0.46%
32.35%
30.98%
0.94%
14.88%
3.50%
15.43%
1.07%
0.37%
0.54%
0.73%
0.15%
1.03%
40.34%
2.25%
1.75%
1.52%
6.33%
6.94%
5.52%
0.83%
14.69%
1.75%
32.11%
0.25%
S2-FITC
hACE2-APC
bACE2$_{R.cor}$-APC
C
hACE2
bACE2$_{R.cor}$-WT
bACE2$_{R.cor}$-ΔGLC$_{38}$
KDa
ACE2-
actin-
-240
-140
-120
-95
-65
-50
-35
D
hACE2
bACE2$_{R.cor}$-WT
bACE2$_{R.cor}$-ΔGLC$_{38}$
RBD$_{SARS2}$-Fc
RBD$_{Rc-o319}$-Fc
Ctr
62.08%
0.07%
0.01%
0.07%
52.26%
0.02%
0.15%
50.95%
0.09%

**Figure S13. Quantification of Rc-o319 S-proteins and SARS-CoV-2 S-protein expression using western blot and flow cytometry.**

**(A)** Total cell S-protein expression was quantified by western blot. Lysates of cells expressing different S-proteins were probed using an anti-FLAG antibody with β-actin as the control. **(B)** Quantification of SARS-CoV-2 and Rc-o319 variant S-protein expression (left panels) and their ACE2 binding (middle panels, hACE2 binding; right panels, bACE2$_{R.cor}$ binding) using flow cytometry. **(C)** Total cell hACE, bACE2$_{R.cor}$ and bACE2$_{R.cor}$-ΔGLC$_{38}$ protein expression was quantified using an anti-flag-tag antibody as the probe by western blot. **(D)** RBD-Fc binding by ACE2 expressing cells as detected by flow cytometry. Cells surface-expressing hACE2, bACE2$_{R.cor}$-WT, or bACE2$_{R.cor}$-ΔGLC$_{38}$ were incubated with SARS-CoV-2 or Rc-o319 RBD-Fc proteins. Binding of dimeric RBD-Fc proteins was quantified using a goat anti-human IgG-APC as the probe.

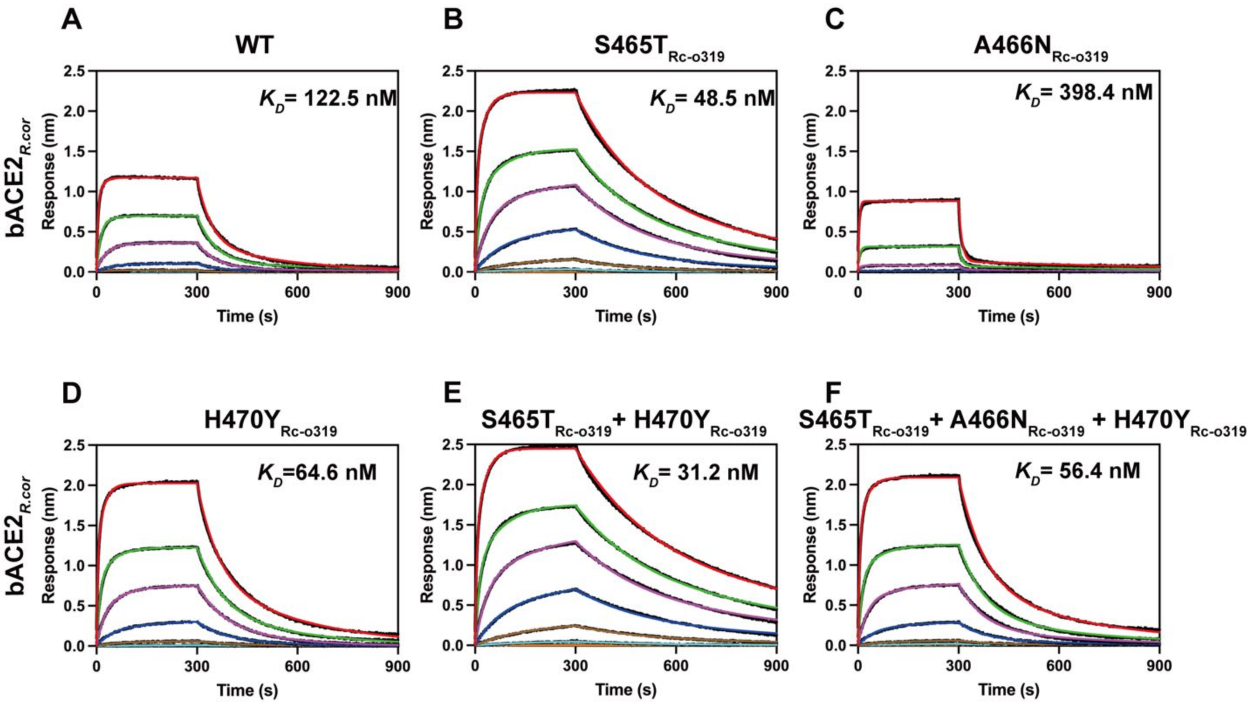


**Figure S14. Binding of bACE2*R.cor* to wild-type (WT) and variant Rc-o319 RBD-Fc proteins carrying individual point mutations in the combined anchor loop (AL) change (S465T$_{Rc\text{-}o319}$+A466N$_{Rc\text{-}o319}$+H470Y$_{Rc\text{-}o319}$) or a double mutation (S465T$_{Rc\text{-}o319}$+H470Y$_{Rc\text{-}o319}$), as measured by BLI assays.**

**(A-F)** Dimeric Rc-o319 RBD-Fc proteins were immobilized on BLI sensors, and dimeric bACE2*R.cor* was used as the analyte in solution, tested using three-fold serial dilutions ranging from 3000 to 4.09 nM. $K_D$ values estimated from each binding experiment are shown alongside the corresponding binding curves.

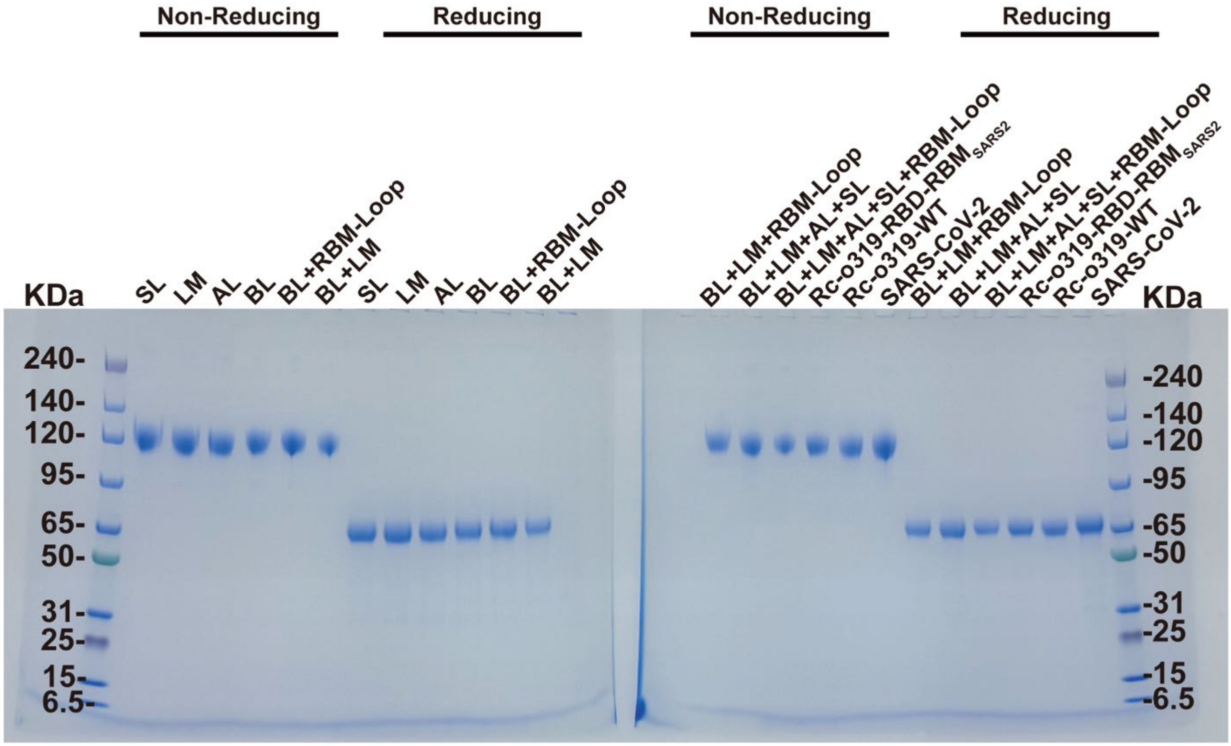


**Figure S15. SDS-PAGE of non-reducing and reducing samples of different Rc-o319 RBD-Fc proteins (10 ug) used for BLI experiments.**

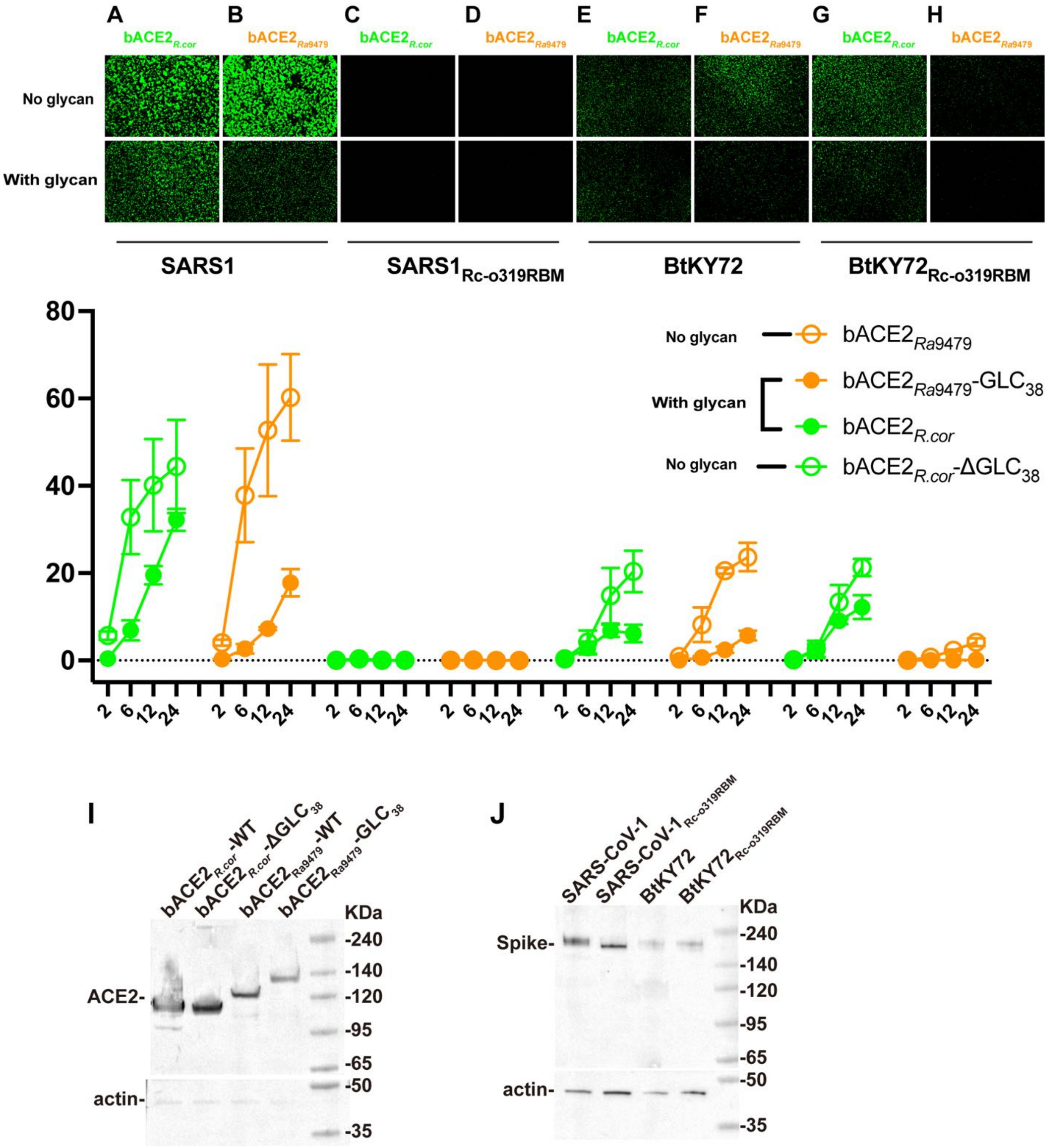


**Figure S16. ACE2 utilization by wild-type (WT) or Rc-o319 RBM chimeric S-proteins (SARS-CoV-1 and BtKY72) in cell-cell fusion assays.**

**(A-H)** The RBMs of BtKY72 and SARS-CoV-1 S-protein were replaced with that of Rc-o319, generating the chimeric $SARS1_{Rc\text{-}o319RBM}$ and $BtKY72_{Rc\text{-}o319RBM}$ S-proteins, respectively. In parallel, we also introduced an Asn38-glycan into $bACE2_{\mathit{Ra}9479}$ by mutation Asp38 to Asn, yielding the $bACE2_{Ra9479}$-$GLC_{38}$

construct. Representative cell-cell fusion images captured at 12 hours post-transfection are shown. Effector cells expressing S-proteins were tested against receptor cells expressing either wild-type or variant of bACE2$_{R.cor}$ or bACE2$_{Ra9479}$. The bottom two panels: Cell-cell fusion was quantified by assessing GFP+ areas at 2, 6, 12 and 24 hours post-transfection. **(I)** Total cell bACE2$_{R.cor}$, bACE2$_{R.cor}$-ΔGLC$_{38}$, bACE2$_{Ra9479}$ and bACE2$_{Ra9479}$-GLC$_{38}$ protein expression was quantified using an anti-flag-tag antibody as the probe by western blot. **(J)** Total cell S-protein expression of SARS-CoV-1, SARS1$_{Rc\text{-}o319RBM}$, BtKY72 and BtKY72$_{Rc\text{-}o319RBM}$ was quantified using an anti-S2-tag antibody as the probe by western blot.

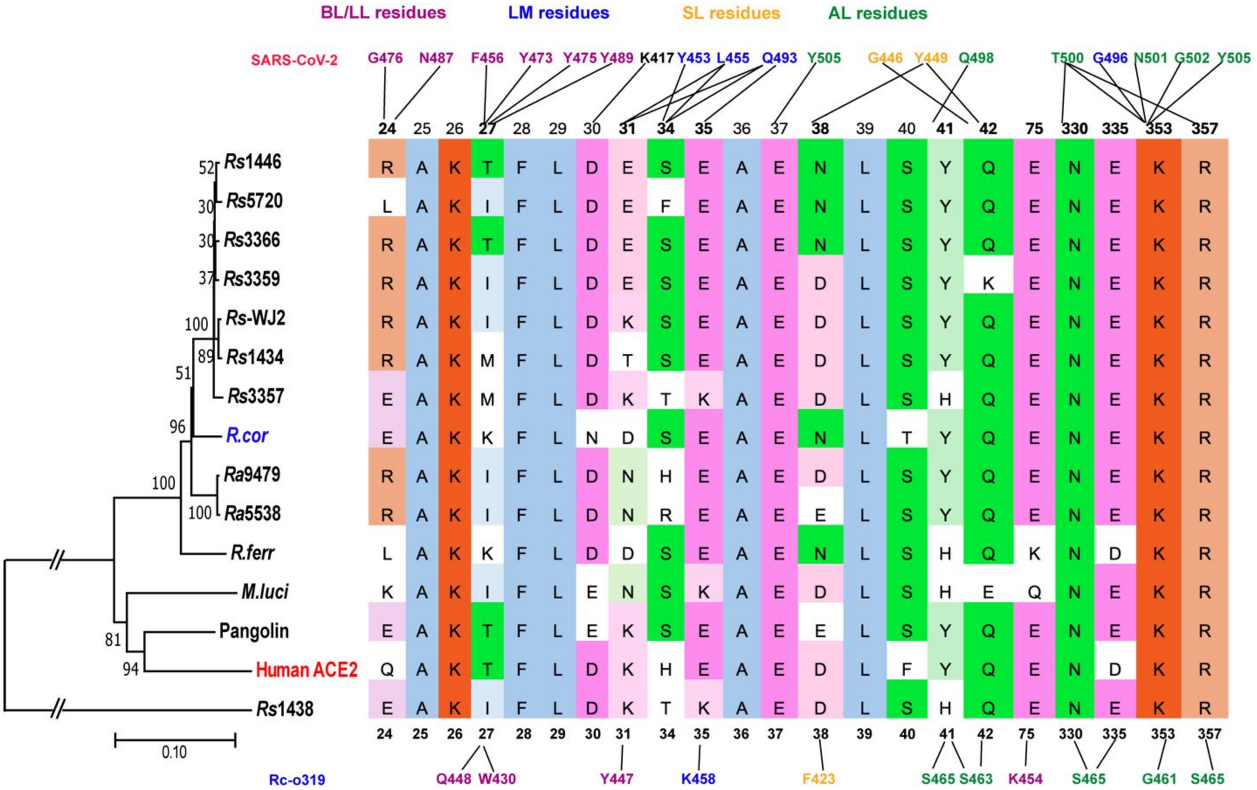


**Figure S17. Phylogenetic tree based on analysis of 15 ACE2 sequences.** The phylogenetic tree was generated with maximum likelihood analysis. The multi-sequence alignment was analyzed using MAFFT and the key interaction residues of sarbecovirus RBD with ACE2 were identified. ACE2 interacting RBM residues of SARS-CoV-2 and Rc-o319 are shown above and below the ACE2 sequence, respectively. Residues in the large loop (LL), lamella (LM), small loop (SL) and anchor loop (AL) regions are colored in purple, blue, orange, and green. Black lines indicate van der Waals contacts, hydrogen bonds, and salt bridges.

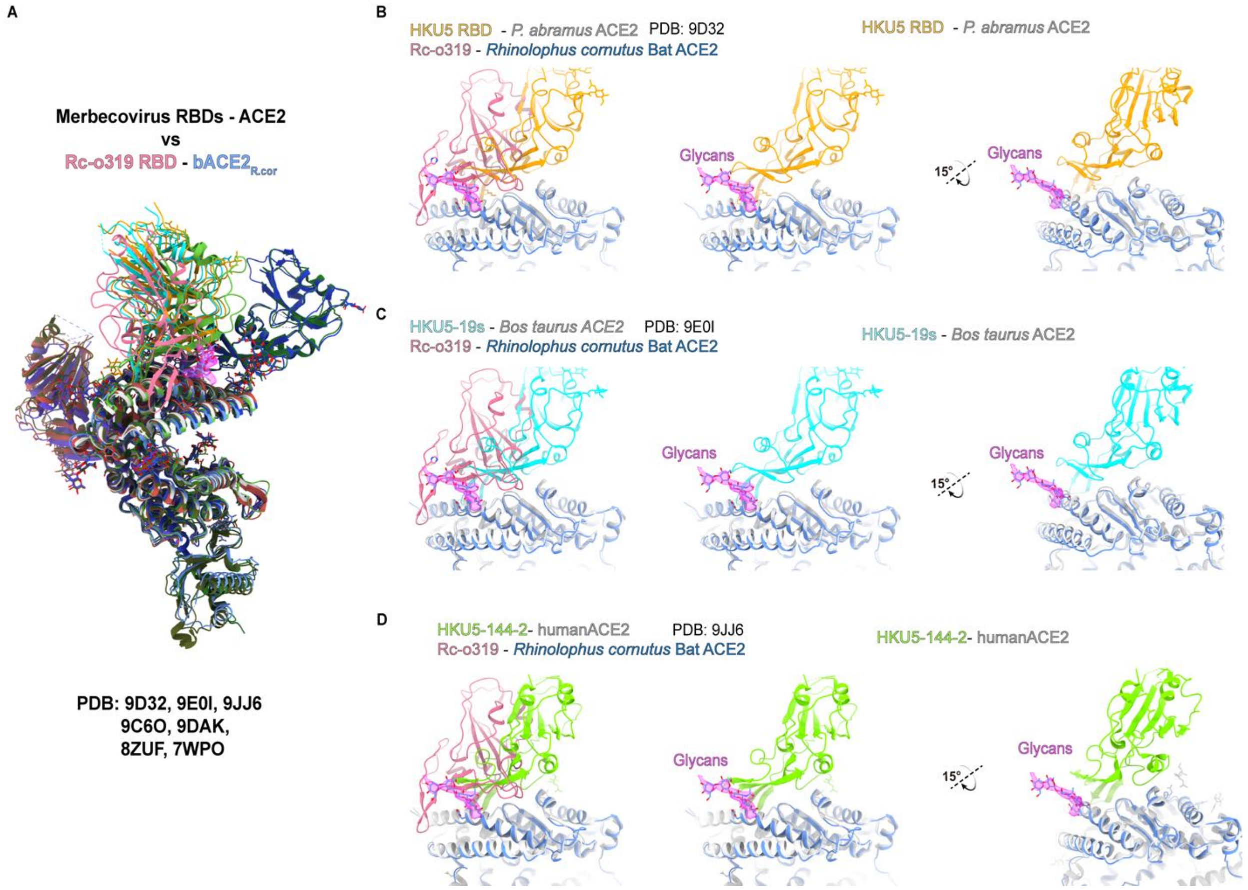


**Figure S18. Comparison of ACE2-bound merbecovirus RBD structures with the Rc-o319 RBD-bACE2*R.cor* complex.**

**(A)** Overview of the structural alignments between the Rc-o319 RBD-bACE2*R.cor* complex and ACE2-bound merbecovirus RBD complexes, including HKU5 RBD-*P.abramus* ACE2 (PDB: 9D32), HKU5-19s-*Bos taurus* ACE2 (PDB: 9E0I), MOW5-22-*P. davyi* ACE2 (PDB: 9C6O), PnNL2018B-*P.nathusii* ACE2 (PDB: 9DAK), MOW15-22-*P.nat* ACE2 (PDB: 8ZUF), NeoCoV-Bat37 ACE2 (PDB: 7WPO), and HKU5-144-2-hACE2 (PDB:9JJ6). **(B–D)** The Asn38-glycan of ACE2 is positioned near the interface between HKU5-like merbecovirus RBDs (HKU5 RBD, HKU5-19s, and HKU5-144-2) and ACE2.

### Table S1. Cryo-EM data collection, refinement and validation statistics.

| | Rc-o319 S-trimer locked-1 conformation | Rc-o319 S-trimer locked-2 conformation | Rc-o319 RBD:bACE2$_{R.cor}$ (C2) | Rc-o319 RBD:bACE2$_{R.cor}$ (focused map) |
|---|---|---|---|---|
| **Data collection and processing** | | | | |
| Magnification | 130000 X | | 130000 X | |
| Voltage (kV) | 300 | | 300 | |
| Electron exposure ($e^{-}$/Å$^2$) | 50 | | 50 | |
| Defocus range (μm) | -0.8 - -2.2 | | -0.8 - -2.0 | |
| Pixel size (Å) | 0.93 | | 0.646 | |
| Movies (no.) | 6255 | | 39866 | |
| Initial particle images (no.) | 732838 | | 1066443 | |
| Symmetry imposed | *C3* | | *C2* | |
| Final particle images (no.) | 86699 | 436097 | 493907 | 987814 |
| Map resolution (Å) | 2.3 | 2.1 | 2.4 | 2.3 |
| FSC threshold | 0.143 | 0.143 | 0.143 | 0.143 |
| Map resolution range (Å) | 2.08-4.33 | 2.08-3.21 | 2.00-3.96 | 1.96-3.97 |
| **Refinement** | | | | |
| Initial model used | 7XU2 | 7XU2 | 8ZY9 | 8ZYA |
| Model resolution (Å) | 3.2 | 3.2 | 2.7 | 2.5 |
| FSC threshold | 0.5 | 0.5 | 0.5 | 0.5 |
| Map sharpening *B* factor (Å$^2$) | 58.1 | 63.8 | 92.5 | 86.1 |
| Model composition | | | | |
| Non-hydrogen atoms | 26247 | 26002 | 14274 | 7315 |
| Protein residues | 3231 | 3182 | 1695 | 868 |
| Ligands | 69 | 75 | 26 | 15 |
| *B* factors (Å$^2$) | | | | |
| Protein | 39.77 | 44.09 | 56.27 | 18.73 |
| Ligand | 79.97 | 56.19 | 103.47 | 35.79 |
| R.m.s. deviations | | | | |
| Bond lengths (Å) | 0.003 | 0.003 | 0.004 | 0.004 |
| Bond angles (°) | 0.575 | 0.652 | 0.781 | 0.698 |
| **Validation** | | | | |
| MolProbity score | 1.23 | 1.63 | 1.75 | 1.71 |
| Clashscore | 2.76 | 3.39 | 5.02 | 4.49 |
| Poor rotamers (%) | 0.00 | 0.00 | 0.00 | 0.00 |
| Ramachandran plot | | | | |
| Favored (%) | 97.64 | 95.94 | 96.36 | 97.10 |
| Allowed (%) | 2.36 | 4.06 | 3.64 | 2.90 |
| Disallowed (%) | 0.00 | 0.00 | 0.00 | 0.00 |

**Table S2. Kinetic parameters of different ACE2 orthologs binding to different constructs of wild-type Rc-o319 S-proteins (related to Figure 1).**

| ACE2 ortholog | Rc-o319-6P S-protein | | | | Rc-o319 S-RBD-Fc | | |
|---|---|---|---|---|---|---|---|
| | $k_{on}$ ($M^{-1}S^{-1}$) | $k_{off}$ ($S^{-1}$) | $K_D$ (nM) | | $k_{on}$ ($M^{-1}S^{-1}$) | $k_{off}$ ($S^{-1}$) | $K_D$ (nM) |
| bACE2$_{R.cor}$ | 9.640x10$^{3}$ ($k_{on1}$) | 2.085x10$^{-2}$ ($k_{off1}$) | 2163 ($k_{off1}/k_{on1}$) | bACE2$_{R.cor}$ | 8.523 x 10$^{4}$ ($k_{on}$) | 6.468 x 10$^{-3}$ ($k_{off}$) | 75.9 ($k_{off}/k_{on}$) |
| | 3.899x10$^{4}$ ($k_{on2}$) | 2.555x10$^{-4}$ ($k_{off2}$) | 534.7 ($k_{off1}/k_{on2}$) | | | | |
| | | | 26.5 ($k_{off2}/k_{on1}$) | | | | |
| | | | 6.6 ($k_{off2}/k_{on2}$) | | | | |
| bACE2$_{Ra9479}$ | - | - | No binding | bACE2$_{Ra9479}$ | - | - | No binding |
| hACE2 | - | - | No binding | hACE2 | - | - | No binding |

### Table S3. Kinetic parameters of different Rc-o319 RBD-Fc variants binding to bACE2$_{R.cor}$ or hACE2 (related to Figure 4).

| Rc-o319 RBD variants | bACE2$_{R.cor}$ | | | Rc-o319 RBD variants | bACE2$_{R.cor}$ | | |
|---|---|---|---|---|---|---|---|
| | $k_{on}$ ($M^{-1}S^{-1}$) | $k_{off}$ ($S^{-1}$) | $K_D$ (nM) | | $k_{on}$ ($M^{-1}S^{-1}$) | $k_{off}$ ($S^{-1}$) | $K_D$ (nM) |
| WT | $8.523 \times 10^4$ ($k_{on}$) | $6.468 \times 10^{-3}$ ($k_{off}$) | 75.9 ($k_{off}/k_{on}$) | BL | - | - | No Binding |
| LM (K458Q) | $1.612 \times 10^4$ ($k_{on}$) | $4.474 \times 10^{-2}$ ($k_{off}$) | 2775 ($k_{off}/k_{on}$) | AL (S465T A466N H470Y) | $6.023 \times 10^4$ ($k_{on}$) | $3.656 \times 10^{-3}$ ($k_{off}$) | 60.7 ($k_{off}/k_{on}$) |
| SL (F423Y) | $8.789 \times 10^4$ ($k_{on}$) | $7.532 \times 10^{-3}$ ($k_{off}$) | 85.7 ($k_{off}/k_{on}$) | BL+ +LM+AL+SL | - | - | No Binding |
| BL+LM+AL+ SL+RBM-loop | - | - | No Binding | Rc-o319-RBD-RBM$_{SARS2}$ | - | - | No Binding |
| BL+LM | - | - | No binding | BL+RBM-loop | $8.215 \times 10^3$ ($k_{on}$) | $3.970 \times 10^{-2}$ ($k_{off}$) | 483 ($k_{off}/k_{on}$) |
| BL+LM+RBM-loop | - | - | No binding | SARS-CoV-2 | - | - | No binding |

| Rc-o319-RBD variants | hACE2 | | | Rc-o319-RBD variants | hACE2 | | |
|---|---|---|---|---|---|---|---|
| | $k_{on}$ ($M^{-1}S^{-1}$) | $k_{off}$ ($S^{-1}$) | $K_D$ (nM) | | $k_{on}$ ($M^{-1}S^{-1}$) | $k_{off}$ ($S^{-1}$) | $K$ (nM) |
| WT | - | - | No Binding | BL | - | - | No Binding |
| LM | - | - | No Binding | AL | - | - | No Binding |
| SL | - | - | No Binding | BL+ LM +AL+SL | - | - | No Binding |
| BL+LM+ AL+SL + RBM-Loop | $2.930 \times 10^4$ ($k_{on}$) | $9.286 \times 10^{-4}$ ($k_{off}$) | 31.7 ($k_{off}/k_{on}$) | Rc-o319-RBD-SARS2$_{RBM}$ | $3.520 \times 10^4$ ($k_{on}$) | $4.331 \times 10^{-4}$ ($k_{off}$) | 12.3 ($k_{off}/k_{on}$) |
| BL+LM | - | - | No binding | BL+RBM-loop | $2.027 \times 10^4$ ($k_{on}$) | $3.218 \times 10^{-2}$ ($k_{off}$) | 1588 ($k_{off}/k_{on}$) |
| BL+LM+RBM-loop | $6.594 \times 10^3$ ($k_{on}$) | $9.155 \times 10^{-3}$ ($k_{off}$) | 1388.4 ($k_{off}/k_{on}$) | SARS-CoV-2 | $7.277 \times 10^3$ ($k_{on}$) | $4.783 \times 10^{-5}$ ($k_{off}$) | 0.065 ($k_{off}/k_{on}$) |

### Table S4. Kinetic parameters of different sarbecovirus RBD-Fc proteins and their variants binding to bACE2$_{R.cor}$ (related to Fig. S9).

| Rc-o319 RBD variants | bACE2$_{R.cor}$ | | | Rc-o319 RBD variants | bACE2$_{R.cor}$ | | |
|---|---|---|---|---|---|---|---|
| | $k_{on}$ ($M^{-1}S^{-1}$) | $k_{off}$ ($S^{-1}$) | $K_D$ (nM) | | $k_{on}$ ($M^{-1}S^{-1}$) | $k_{off}$ ($S^{-1}$) | $K_D$ (nM) |
| WT | $6.863 \times 10^4$ ($k_{on}$) | $1.654 \times 10^{-2}$ ($k_{off}$) | 241.0 ($k_{off}/k_{on}$) | K454Y | $1.476 \times 10^5$ ($k_{on}$) | $1.803 \times 10^{-1}$ ($k_{off}$) | 1221.5 ($k_{off}/k_{on}$) |
| LM (K458Q) | $1.612 \times 10^4$ ($k_{on}$) | $4.474 \times 10^{-2}$ ($k_{off}$) | 2775 ($k_{off}/k_{on}$) | LM (K458A) | $6.177 \times 10^4$ ($k_{on}$) | $1.483 \times 10^{-1}$ ($k_{off}$) | Weak binding |
| Sarbecovirus-RBD variants | bACE2$_{R.cor}$ | | | Sarbecovirus-RBD variants | bACE2$_{R.cor}$ | | |
| | $k_{on}$ ($M^{-1}S^{-1}$) | $k_{off}$ ($S^{-1}$) | $K_D$ (nM) | | $k_{on}$ ($M^{-1}S^{-1}$) | $k_{off}$ ($S^{-1}$) | $K_D$ (nM) |
| BANAL-20-52 | $9.891 \times 10^4$ ($k_{on}$) | $6.288 \times 10^{-2}$ ($k_{off}$) | Weak binding ($k_{off}/k_{on}$) | BANAL-20-236 | $2.731 \times 10^4$ ($k_{on}$) | $3.580 \times 10^{-4}$ ($k_{off}$) | 13.1 ($k_{off}/k_{on}$) |
| SARS1-N479K | - | - | Weak binding | SARS2-Q493K | $3.191 \times 10^4$ ($k_{on}$) | $2.435 \times 10^{-2}$ ($k_{off}$) | 763.0 ($k_{off}/k_{on}$) |

**Table S5. Kinetic parameters of hACE2, bACE2$_{R.cor}$ or bACE2$_{R.cor}$-ΔGLC$_{38}$ binding to different Rc-o319 S-trimers (related to Fig. S10).**

| Spike | bACE2$_{R.cor}$-WT | | | hACE2 | | |
|---|---|---|---|---|---|---|
| | $k_{on}$ ($M^{-1}S^{-1}$) | $k_{off}$ ($S^{-1}$) | $K_D$ (nM) | $k_{on}$ ($M^{-1}S^{-1}$) | $k_{off}$ ($S^{-1}$) | $K_D$ (nM) |
| Rc-o319 | $7.746 \times 10^{3}$ ($k_{on1}$)<br>$1.045 \times 10^{5}$ ($k_{on2}$) | $1.467 \times 10^{-2}$ ($k_{off1}$)<br>$4.303 \times 10^{-4}$ ($k_{off2}$) | 1894 ($k_{off1}/k_{on1}$)<br>140.4 ($k_{off1}/k_{on2}$)<br>55.6 ($k_{off2}/k_{on1}$)<br>4.1 ($k_{off2}/k_{on2}$) | - | - | No Binding |
| | $k_{on}$ ($M^{-1}S^{-1}$) | $k_{off}$ ($S^{-1}$) | $K_D$ (nM) | $k_{on}$ ($M^{-1}S^{-1}$) | $k_{off}$ ($S^{-1}$) | $K_D$ (nM) |
| BL | - | - | No Binding | $2.966 \times 10^{3}$ ($k_{on1}$)<br>$7.408 \times 10^{4}$ ($k_{on2}$) | $9.203 \times 10^{-2}$ ($k_{off1}$)<br>$1.193 \times 10^{-3}$ ($k_{off2}$) | 31024 ($k_{off1}/k_{on1}$)<br>1242.3 ($k_{off1}/k_{on2}$)<br>402.0 ($k_{off2}/k_{on1}$)<br>16.1 ($k_{off2}/k_{on2}$) |
| | $k_{on}$ ($M^{-1}S^{-1}$) | $k_{off}$ ($S^{-1}$) | $K_D$ (nM) | $k_{on}$ ($M^{-1}S^{-1}$) | $k_{off}$ ($S^{-1}$) | $K_D$ (nM) |
| AL | $4.557 \times 10^{3}$ ($k_{on1}$)<br>$4.437 \times 10^{4}$ ($k_{on2}$) | $1.333 \times 10^{-2}$ ($k_{off1}$)<br>$2.223 \times 10^{-4}$ ($k_{off2}$) | 2925.0 ($k_{off1}/k_{on1}$)<br>300.4 ($k_{off1}/k_{on2}$)<br>48.8 ($k_{off2}/k_{on1}$)<br>5.0 ($k_{off2}/k_{on2}$) | - | - | No Binding |
| | $k_{on}$ ($M^{-1}S^{-1}$) | $k_{off}$ ($S^{-1}$) | $K_D$ (nM) | $k_{on}$ ($M^{-1}S^{-1}$) | $k_{off}$ ($S^{-1}$) | $K_D$ (nM) |
| BL+LM+AL +SL | - | - | No Binding | $5.649 \times 10^{3}$ ($k_{on1}$)<br>$1.621 \times 10^{5}$ ($k_{on2}$) | $5.611 \times 10^{-2}$ ($k_{off1}$)<br>$6.281 \times 10^{-4}$ ($k_{off2}$) | 9933.5 ($k_{off1}/k_{on1}$)<br>346.2 ($k_{off1}/k_{on2}$)<br>111.2 ($k_{off2}/k_{on1}$)<br>3.9 ($k_{off2}/k_{on2}$) |

| | | | | | | |
|---|---|---|---|---|---|---|
| | $k_{on}$ ($M^{-1}S^{-1}$) | $k_{off}$ ($S^{-1}$) | $K_D$ (nM) | $k_{on}$ ($M^{-1}S^{-1}$) | $k_{off}$ ($S^{-1}$) | $K_D$ (nM) |
| BL+LM+AL +SL +RBM-loop | - | - | No Binding | $1.948 \times 10^{3}$ ($k_{on1}$) | $3.316 \times 10^{-4}$ ($k_{off1}$) | 170.3 ($k_{off1}/k_{on1}$) |
| | | | | $1.211 \times 10^{4}$ ($k_{on2}$) | $< 1 \times 10^{-7}$ ($k_{off2}$) | 27.4 ($k_{off1}/k_{on2}$) |
| | | | | | | < 0.01 ($k_{off2}/k_{on1}$) |
| | | | | | | < 0.01 ($k_{off2}/k_{on2}$) |
| ACE2 | | BL | | | BL+LM+AL+SL +RBM-loop | |
| | $k_{on}$ ($M^{-1}S^{-1}$) | $k_{off}$ ($S^{-1}$) | $K_D$ (nM) | $k_{on}$ ($M^{-1}S^{-1}$) | $k_{off}$ ($S^{-1}$) | $K_D$ (nM) |
| bACE2$_{R.cor}$-$\Delta$GLC$_{38}$ | $8.317 \times 10^{3}$ ($k_{on1}$) | $5.669 \times 10^{-2}$ ($k_{off1}$) | 6816.2 ($k_{off1}/k_{on1}$) | $4.774 \times 10^{3}$ ($k_{on1}$) | $9.353 \times 10^{-2}$ ($k_{off1}$) | 19593 ($k_{off1}/k_{on1}$) |
| | $9.335 \times 10^{7}$ ($k_{on2}$) | $1.110 \times 10^{-3}$ ($k_{off2}$) | 133.4 ($k_{off1}/k_{on2}$) | $2.253 \times 10^{5}$ ($k_{on2}$) | $< 1 \times 10^{-7}$ ($k_{off2}$) | 415.2 ($k_{off1}$/k$_{on2}$) |
| | | | 0.6 ($k_{off2}/k_{on1}$) | | | <0.01 ($k_{off2}/k_{on1}$) |
| | | | 0.01 ($k_{off2}/k_{on2}$) | | | <0.01 ($k_{off2}/k_{on2}$) |

### Table S6. Kinetic parameters of bACE2$_{R.cor}$ or bACE2$_{R.cor}$-ΔGLC$_{38}$ binding to different sarbecovirus S-trimers (related to Fig. S11A-H).

| Spike | bACE2$_{R.cor}$-WT | | | bACE2$_{R.cor}$-ΔGLC$_{38}$ | | |
|---|---|---|---|---|---|---|
| | $k_{on}$ ($M^{-1}S^{-1}$) | $k_{off}$ ($S^{-1}$) | $K_D$ (nM) | $k_{on}$ ($M^{-1}S^{-1}$) | $k_{off}$ ($S^{-1}$) | $K_D$ (nM) |
| Rc-o319 | $3.131 \times 10^3$ ($k_{on1}$)<br>$3.503 \times 10^4$ ($k_{on2}$) | $1.048 \times 10^{-2}$ ($k_{off1}$)<br>$3.474 \times 10^{-4}$ ($k_{off2}$) | 3347.5 ($k_{off1}/k_{on1}$)<br>299.2 ($k_{off1}/k_{on2}$)<br>111.0 ($k_{off2}/k_{on1}$)<br>9.9 ($k_{off2}/k_{on2}$) | $5.372 \times 10^3$ ($k_{on1}$)<br>$2.585 \times 10^4$ ($k_{on2}$) | $2.347 \times 10^{-3}$ ($k_{off1}$)<br>$< 1 \times 10^{-7}$ ($k_{off2}$) | 436.9 ($k_{off1}/k_{on1}$)<br>90.8 ($k_{off1}/k_{on2}$)<br><0.0186 ($k_{off2}/k_{on1}$)<br><0.0038 ($k_{off2}/k_{on2}$) |
| | $k_{on}$ ($M^{-1}S^{-1}$) | $k_{off}$ ($S^{-1}$) | $K_D$ (nM) | $k_{on}$ ($M^{-1}S^{-1}$) | $k_{off}$ ($S^{-1}$) | $K_D$ (nM) |
| BANAL-20-52 | - | - | No Binding | $3.049 \times 10^3$ ($k_{on1}$)<br>$3.109 \times 10^3$ ($k_{on2}$) | $6.073 \times 10^{-5}$ ($k_{off1}$)<br>$< 1 \times 10^{-7}$ ($k_{off2}$) | 19.9 ($k_{off1}/k_{on1}$)<br>19.6 ($k_{off1}/k_{on2}$)<br><0.033 ($k_{off2}/k_{on1}$)<br><0.032 ($k_{off2}/k_{on2}$) |
| BANAL-20-236 | $6.635 \times 10^3$ ($k_{on1}$)<br>$7.280 \times 10^3$ ($k_{on2}$) | $2.194 \times 10^{-4}$ ($k_{off1}$)<br>$<1 \times 10^{-7}$ ($k_{off2}$) | 33.1 ($k_{off1}/k_{on1}$)<br>30.1 ($k_{off1}/k_{on2}$)<br><0.015 ($k_{off2}/k_{on1}$)<br><0.014 ($k_{off2}/k_{on2}$) | $6.800 \times 10^3$ ($k_{on1}$)<br>$3.320 \times 10^4$ ($k_{on2}$) | $2.234 \times 10^{-4}$ ($k_{off1}$)<br>$<1 \times 10^{-7}$ ($k_{off2}$) | 32.9 ($k_{off1}/k_{on1}$)<br>6.73 ($k_{off1}/k_{on2}$)<br><0.015 ($k_{off2}/k_{on1}$)<br><0.003 ($k_{off2}/k_{on2}$) |
| | $k_{on}$ ($M^{-1}S^{-1}$) | $k_{off}$ ($S^{-1}$) | $K_D$ (nM) | $k_{on}$ ($M^{-1}S^{-1}$) | $k_{off}$ ($S^{-1}$) | $K_D$ (nM) |
| BtKY72 | - | - | No Binding | $5.637 \times 10^3$ ($k_{on1}$)<br>$2.747 \times 10^4$ ($k_{on2}$) | $1.016 \times 10^{-2}$ ($k_{off1}$)<br>$3.250 \times 10^{-4}$ ($k_{off2}$) | 1802.7 ($k_{off1}/k_{on1}$)<br>370.0 ($k_{off1}/k_{on2}$)<br>57.7 ($k_{off2}/k_{on1}$)<br>11.8 ($k_{off2}/k_{on2}$) |

**Table S7. Kinetic parameters of bACE2$_{Ra9479}$-GLC$_{38}$ or bACE2$_{Ra9479}$ binding to different sarbecovirus S-trimers (related to Fig. S11J and K).**

| Spike | bACE2$_{Ra9479}$-GLC$_{38}$ | | | bACE2$_{Ra9479}$-WT | | |
|---|---|---|---|---|---|---|
| | $k_{on}$ ($M^{-1}S^{-1}$) | $k_{off}$ ($S^{-1}$) | $K_D$ (nM) | $k_{on}$ ($M^{-1}S^{-1}$) | $k_{off}$ ($S^{-1}$) | $K_D$ (nM) |
| SARS-CoV-1 | 3.995 x $10^3$ ($k_{on1}$) | 1.886 x $10^{-2}$ ($k_{off1}$) | 4722.4 ($k_{off1}/k_{on1}$) | 4.073 x $10^3$ ($k_{on1}$) | 2.290 x $10^{-4}$ ($k_{off1}$) | 56.2 ($k_{off1}/k_{on1}$) |
| | 7.279 x $10^3$ ($k_{on2}$) | 1.152 x $10^{-4}$ ($k_{off2}$) | 2591.6 ($k_{off1}/k_{on2}$) | 3.079 x $10^4$ ($k_{on2}$) | 3.784 x $10^{-7}$ ($k_{off2}$) | 7.4 ($k_{off1}/k_{on2}$) |
| | | | 28.8 ($k_{off2}/k_{on1}$) | | | 0.093 ($k_{off2}/k_{on1}$) |
| | | | 15.8 ($k_{off2}/k_{on2}$) | | | 0.012 ($k_{off2}/k_{on2}$) |
| | $k_{on}$ ($M^{-1}S^{-1}$) | $k_{off}$ ($S^{-1}$) | $K_D$ (nM) | $k_{on}$ ($M^{-1}S^{-1}$) | $k_{off}$ ($S^{-1}$) | $K_D$ (nM) |
| BtKY72 | 8.959 x $10^3$ ($k_{on1}$) | 1.234 x $10^{-2}$ ($k_{off1}$) | 1377.8 ($k_{off1}/k_{on1}$) | 4.134 x $10^3$ ($k_{on1}$) | 2.182x $10^{-3}$ ($k_{off1}$) | 528.0 ($k_{off1}/k_{on1}$) |
| | 9.159 x $10^4$ ($k_{on2}$) | 1.909 x $10^{-4}$ ($k_{off2}$) | 134.7 ($k_{off1}/k_{on2}$) | 5.934x $10^4$ ($k_{on2}$) | < 1 x $10^{-7}$ ($k_{off2}$) | 36.8 ($k_{off1}/k_{on2}$) |
| | | | 21.3 ($k_{off2}/k_{on1}$) | | | 0.012 ($k_{off2}/k_{on1}$) |
| | | | 2.1 ($k_{off2}/k_{on2}$) | | | <0.01 ($k_{off2}/k_{on2}$) |

**Table S8. Kinetic parameters of different variants of SARS-CoV-2-RBD-Fc protein binding to bACE2$_{\textit{R.cor}}$ or hACE2 (related to Fig. S12).**

| SARS-CoV-2-RBD-Fc variants | bACE2$_{R.cor}$ | | | hACE2 | | |
|---|---|---|---|---|---|---|
| | $k_{on}$ ($M^{-1}S^{-1}$) | $k_{off}$ ($S^{-1}$) | $K_D$ (nM) | $k_{on}$ ($M^{-1}S^{-1}$) | $k_{off}$ ($S^{-1}$) | $K_D$ (nM) |
| $BL_{Rc-o319}$ | - | - | No binding | - | - | No binding |
| $BL_{Rc-o319}$ + RBM-loop$_{Rc-o319}$ | - | - | No binding | - | - | No binding |
| $BL_{Rc-o319}$ + $LM_{Rc-o319}$ | - | - | Weak binding | - | - | No binding |
| $RBM_{Rc-o319}$ | 8.133x $10^4$ ($k_{on}$) | 6.596 x $10^{-3}$ ($k_{off}$) | 81.1 ($k_{off}/k_{on}$) | - | - | No binding |

**Table S9. Kinetic parameters of different Rc-o319 RBD-Fc variants binding to bACE2$_{R.cor}$ (related to Fig. S14).**

| Rc-o319 RBD variants | bACE2$_{R.cor}$ | | | Rc-o319 RBD variants | bACE2$_{R.cor}$ | | |
|---|---|---|---|---|---|---|---|
| | $k_{on}$ (M$^{-1}$S$^{-1}$) | $k_{off}$ (S$^{-1}$) | $K_D$ (nM) | | $k_{on}$ (M$^{-1}$S$^{-1}$) | $k_{off}$ (S$^{-1}$) | $K_D$ (nM) |
| WT | 2.385 x 10$^{4}$ ($k_{on}$) | 2.922 x 10$^{-3}$ ($k_{off}$) | 122.5 ($k_{off}$/$k_{on}$) | S465T | 3.510 x 10$^{4}$ ($k_{on}$) | 1.701 x 10$^{-3}$ ($k_{off}$) | 48.5 ($k_{off}$/$k_{on}$) |
| A466N | 1.147 x 10$^{5}$ ($k_{on}$) | 4.568 x 10$^{-2}$ ($k_{off}$) | 398.4 ($k_{off}$/$k_{on}$) | H470Y | 4.621 x 10$^{4}$ ($k_{on}$) | 2.987 x 10$^{-3}$ ($k_{off}$) | 64.6 ($k_{off}$/$k_{on}$) |
| S465T +A466N | 3.454 x 10$^{4}$ ($k_{on}$) | 1.077 x10$^{-3}$ ($k_{off}$) | 31.2 ($k_{off}$/$k_{on}$) | S465T+ A466N+ H470Y | 4.429 x 10$^{4}$ ($k_{on}$) | 2.498 x10$^{-3}$ ($k_{off}$) | 56.4 ($k_{off}$/$k_{on}$) |

### Table S10. Key ACE2 residues in the ACE2-RBD interface of the orthologs tested in the cell-cell fusion assay (related to Figure 1 and supplymentary Fig.17).

| Species | Isolate # | Genbank | ACE2-RBD Contact Residues | | | | | | | | | | | | | | | |
|---|---|---|---|---|---|---|---|---|---|---|---|---|---|---|---|---|---|---|
| | | | **24** | **27** | **30** | **31** | **34** | **35** | **38** | **40** | **41** | **42** | **45** | **82** | **83** | **330** | **353** | **355** |
| ***R. cornutus*** | *R.cor* | BCG67443.1 | E | **K** | N | D | S | E | **N** | T | Y | Q | L | N | Y | N | K | D |
| ***R. sinicus*** | WJ2 | QMQ39202.1 | R | **I** | D | K | S | E | D | S | Y | Q | L | N | Y | N | K | D |
| ***R. sinicus*** | 1446 | MT394194.1 | R | **T** | D | E | S | E | N | S | Y | Q | L | N | Y | N | K | D |
| ***R. sinicus*** | 1434 | QMQ39216.1 | R | **M** | D | T | S | E | D | S | Y | Q | L | N | Y | N | K | D |
| ***R. sinicus*** | 1438 | MT394184.1 | E | **I** | D | K | T | K | D | S | H | Q | L | N | Y | N | K | D |
| ***R. sinicus*** | 3357 | AGZ48803.1 | E | **M** | D | K | T | K | D | S | H | Q | L | N | Y | N | K | D |
| ***R. sinicus*** | 3366 | QMQ39215.1 | R | **T** | D | E | S | E | **N** | S | Y | Q | L | N | Y | N | K | D |
| ***R. sinicus*** | 3359 | QMQ39211.1 | R | **I** | D | E | S | E | D | S | Y | K | L | N | Y | N | K | D |
| ***R. sinicus*** | 5720 | MT394182.1 | L | **I** | D | E | F | E | **N** | S | Y | Q | L | N | Y | N | K | D |
| ***R. affinis*** | 9479 | MT394208.1 | R | **I** | D | N | H | E | D | S | Y | Q | L | N | Y | N | K | D |
| ***R. affinis*** | 787 | MT394203.1 | R | **I** | D | N | R | E | E | S | Y | Q | L | N | Y | N | K | D |
| ***R. ferrumequinum*** | *R.ferr* | BAH02663.1 | L | **K** | D | D | S | E | **N** | S | H | Q | L | N | F | N | K | D |
| ***Myotis lucifugus*** | *M.luci* | XP_023609438.1 | K | **I** | E | N | S | K | D | S | H | E | L | N | Y | N | K | D |
| **Pangolin** | - | XP_017505752.1 | E | **T** | E | K | S | E | E | S | Y | Q | L | N | Y | N | K | D |
| **Human** | - | BAB40370.1 | Q | **T** | D | K | H | E | D | F | Y | Q | L | M | Y | N | K | D |
